# Glucocorticoids impair memory inference through noradrenergic disruption of GABAergic regulation in hippocampal CA3

**DOI:** 10.64898/2026.09.02.748771

**Authors:** Emma Clephas, Bente Raaso, Sara Spolaore, Lars Schwabe, Benno Roozendaal

## Abstract

The integration of separate memories sharing overlapping elements allows for inferring relationships between events that have never been experienced together. Recent findings indicate that stress impairs this process of mnemonic inference, which may have important implications for stress-related psychopathologies. However, the mechanisms underlying such impairments remain largely unknown. To elucidate the neuroendocrine mechanisms by which stress affects mnemonic inference, mice were first trained on two sets of auditory-visual cue associations. Afterwards, one of the visual cues was paired with reward. Mnemonic inference was probed by examining reward-seeking behavior in response to the auditory cues alone. Control mice successfully inferred the auditory cue-outcome association, selectively seeking reward in response to the auditory cue associated with the rewarded visual cue. In contrast, systemic administration of the stress hormone corticosterone before testing impaired inference accuracy, leading mice to seek reward to both auditory cues. Corticosterone administration was accompanied by increased noradrenergic activity, and stimulation of noradrenergic signaling with yohimbine similarly disrupted inference accuracy. Conversely, the β-adrenoceptor antagonist propranolol prevented the corticosterone-induced impairment, indicating that corticosterone effects depend on concurrent noradrenergic activation. At the neural level, the corticosterone-induced mnemonic inference impairment was associated with enhanced GABAergic inhibition of pyramidal neurons in the hippocampal CA3 region. Furthermore, hippocampal administration of the GABA_A_ receptor antagonist bicuculline prevented the corticosterone-induced impairment in mnemonic inference. Together, these findings identify a glucocorticoid–noradrenergic mechanism that disrupts mnemonic inference through enhanced GABAergic inhibition in hippocampal CA3, providing an account of how stress hormones impair inferential memory.

## Introduction

Stressful or emotionally arousing experiences leave lasting memories through the actions of glucocorticoids (corticosterone in rodents; CORT) and norepinephrine^1^. Both stress hormones interact to enhance memory consolidation and impair memory retrieval^1–3^. Further, evidence shows that stress hormones can induce a shift from flexible, cognitive memory systems such as the hippocampus, to more habitual systems such as the dorsal striatum^4–7^, impacting the flexibility of memory. A key aspect of mnemonic flexibility is integrating overlapping experiences to infer relationships between events that were never directly experienced together^8,9^. Recent work in humans indicated that stress exposure can impair inference^10,11^. Such stress-induced inference deficits might have major implications for, e.g., eyewitness testimony or stress-related psychopathologies. Yet, the underlying mechanisms are largely unknown.

Successful memory inference relies on the hippocampus, and its interaction with the medial prefrontal cortex^9^. The hippocampus plays a crucial role in the consolidation, retrieval, and subsequent integration of memories^8,12^. During recall, the hippocampus supports the formation of initial links between related experiences. Especially hippocampal area CA3 has been associated with these initial links^13,14^. The hippocampus is highly sensitive to stress^15–17^. It expresses abundant glucocorticoid receptors^18,19^, and CORT strongly modulates hippocampal function^20^. Evidence further indicates that these glucocorticoid effects depend, at least partly, on concurrent noradrenergic signaling^21–23^. At the neural level, CORT and norepinephrine can alter circuit function through modulation of glutamatergic^24,25^ and GABAergic transmission^26,27^. Together, these findings suggest that stress hormones influence mnemonic flexibility by altering the balance of excitatory and inhibitory signaling within the hippocampal circuit. However, whether CORT affects memory inference through such mechanisms, and to what extent these effects depend on noradrenergic signaling, remains unknown.

Here, we tested whether CORT shapes hippocampal contributions to mnemonic inference. Mice were trained on a multi-day memory inference task^8^, in which they learned overlapping auditory– visual cue associations. Subsequently, one visual cue was paired with reward, and, following systemic CORT administration, inference was tested by re-exposure to the auditory cue.

Additionally, we investigated interactions between CORT and norepinephrine on memory inference. We further assessed the involvement of the different hippocampal subregions in mediating the CORT effects on memory inference by examining activity of both GABAergic and glutamatergic cells during the memory inference test. As we found that CORT administration increased GABAergic inhibition of hippocampal pyramidal cell activity, we next administered CORT and the GABA_A_ receptor antagonist bicuculline directly into the hippocampus. We hypothesized that CORT, acting through noradrenergic and GABAergic mechanisms, alters hippocampal activity and, as such, impairs mnemonic inference.

## Materials and Methods

### Subjects

Male C57BL/6J mice (8-10 weeks; Charles River; France) were single housed under a standard 12-h/12-h light/dark cycle, with *ad libitum* food access until 2 days before conditioning, after which they were maintained at 90% of free-feeding body weight. Training and testing were performed during the light phase between 09:00 and 16:00 h. All procedures complied with EU directive 2010/63/EU and were approved by the Central Authority for Scientific Procedures on Animals, The Hague, The Netherlands.

### Memory inference task

Before training, mice were handled for 4 min per day for 3 days. Next, animals were trained and tested on a memory inference task, consisting of observational learning, conditioning and inference testing phases^8^. During observational learning, animals learned auditory–visual cue associations (2 kHz tone–orange circle; 500 Hz tone–green L-shape). During conditioning, one visual cue was paired with a 15% sucrose reward. During testing, auditory cues were presented alone and reward-seeking behavior was measured.

Behavior was video-recorded and scored offline by an observer blind to the treatment. A reward-seeking bias (i.e., [reward-seeking behavior rewarded – reward-seeking behavior non-rewarded]/[total reward-seeking behavior] × 100 %) was computed. Additional control experiments and detailed descriptions of the task, equipment and analysis procedures are provided in the Supplementary Methods.

### Systemic drug administration

CORT (3 or 10 mg/kg; Sigma-Aldrich) was dissolved in 5% ethanol in saline^28^. Animals received CORT, CORT and (±)-propranolol hydrochloride (1 mg/kg; Sigma-Aldrich), vehicle (5% ethanol in saline) or propranolol. The noradrenergic stimulant yohimbine (1 or 3 mg/kg; Sigma-Aldrich) was dissolved in saline^29,30^. Control animals received saline. All treatments were administered intraperitoneally (10 mL/kg) 45 min prior to the memory inference test. Drug solutions were prepared freshly before each experiment.

### Local corticosterone administration

CORT (3 ng; Sigma-Aldrich) was dissolved in 0.25% ethanol in saline (vehicle). Animals received CORT, CORT and the GABA_A_ receptor antagonist bicuculline methiodide (0.5 ng; ThermoFisher), vehicle or bicuculline. Drugs were administered directly into the hippocampus through bilateral cannulas using a 1-μL Hamilton microsyringe (30-gauge needle, 7002 series) in a volume of 0.3 μL per hemisphere 45 min prior to the memory inference test.

### Surgery for cannula implantation

Mice were anesthetized with isoflurane (5.0% for induction, 1.0-2.0% for maintenance). Perioperative analgesia was provided with carprofen in the drinking water, and a local 2% lidocaine-bupivacaine solution. Bilateral guide cannulas (Bio Services #6100050, diameter: 0.48 mm, length: 2.5 mm, 23 gauge) were inserted into the dorsal hippocampus (anteroposterior (AP): - 1.7 mm, relative to Bregma, mediolateral (ML): ±1.2 mm, dorsoventral (DV): -1.9 mm)^31^, and secured with dental cement (Super-Bond Universal Kit #812246; Dental Bauer). Animals were allowed to recover for >7 days.

### Neuronal activity measurements

Thirty minutes after the memory inference test, animals were killed by cervical dislocation, the brains rapidly dissected, and frozen in isopentane for use in fluorescent in situ hybridization (FISH) protocols. Dorsal hippocampal sections (AP: -1.46 mm to -2.30 mm) were analyzed for *c-Fos* expression in glutamatergic (*Slc17a7*-positive) and GABAergic (*Slc32a1*-positive) neurons.

In separate cohorts, mice were anesthetized with an overdose of sodium pentobarbital 90 min after the memory inference test, followed by transcardial perfusion with 0.1 M phosphate-buffered saline (PBS) and 4% paraformaldehyde (PFA, pH 7.2). Brains were post-fixated in 4% PFA at 4 °C overnight. Dorsal hippocampal sections (AP: -1.46 mm to -2.30 mm) were analyzed for c-Fos expression in GABAergic (GAD67-positive) neurons. Additional staining assessed phosphorylated GABA_A_ receptor expression in pyramidal neurons. Finally, to evaluate noradrenergic activity, c-Fos expression was quantified in tyrosine hydroxylase-positive neurons of the locus coeruleus (LC; AP: -5.34 mm to -5.68 mm). Detailed staining protocols and antibody information are provided in the Supplementary Methods.

### Imaging and quantification

FISH and immunohistochemistry sections were imaged using the Axioobserver with Sample Finder AI from Zeiss at 20X magnification. Analyses focused on the dentate gyrus, CA1 and CA3 subregions of the dorsal hippocampus. Fluorescent cells were quantified using an automated pipeline for FISH and blind manual counting for immunohistochemistry, separately for each subregion. Quantifications were normalized to the area (cells/mm^2^), and averaged across four sections per animal. Detailed image acquisition and analysis procedures are provided in the Supplementary Methods.

### Corticosterone Assay

Thirty minutes after the memory inference test, trunk blood was collected in lithium heparin-coated tubes (Sarstedt) and kept on ice until centrifugation (10,000 rpm for 10 min). Plasma CORT concentration levels were measured in duplicate with the Enzyme Linked Immune Sorbent Assay (ELISA) Corticosterone kit (Enzo Life Sciences; ADI-901-097) according to the manufacturer’s protocol.

### Statistical analyses

All statistical analyses were performed using JASP 0.16.3.0.

The reward-seeking bias on the memory inference test was analyzed with one- or two-way or repeated measures ANOVA. For FISH and immunohistochemistry, the number of fluorescent neurons was analyzed with independent samples *t*-tests or two-way ANOVA. Plasma CORT levels were log-transformed and one-way ANOVA was used to analyze differences between drug treatment groups.

When appropriate, post-hoc tests were used to determine the origin of the significance, and *P-* values were corrected for multiple testing using a Bonferroni correction. All statistical tests were two-tailed, and *P* < 0.05 was accepted as statistically significant. Detailed descriptions of the statistical analyses are provided in the Supplementary Methods.

## Results

### Systemic CORT administration impairs mnemonic inference

To examine whether systemic CORT administration affects memory inference, mice were trained on a memory inference task, and received CORT (3 or 10 mg/kg) or vehicle 45 min before the inference test (Figure 1A).

**Figure 1.**
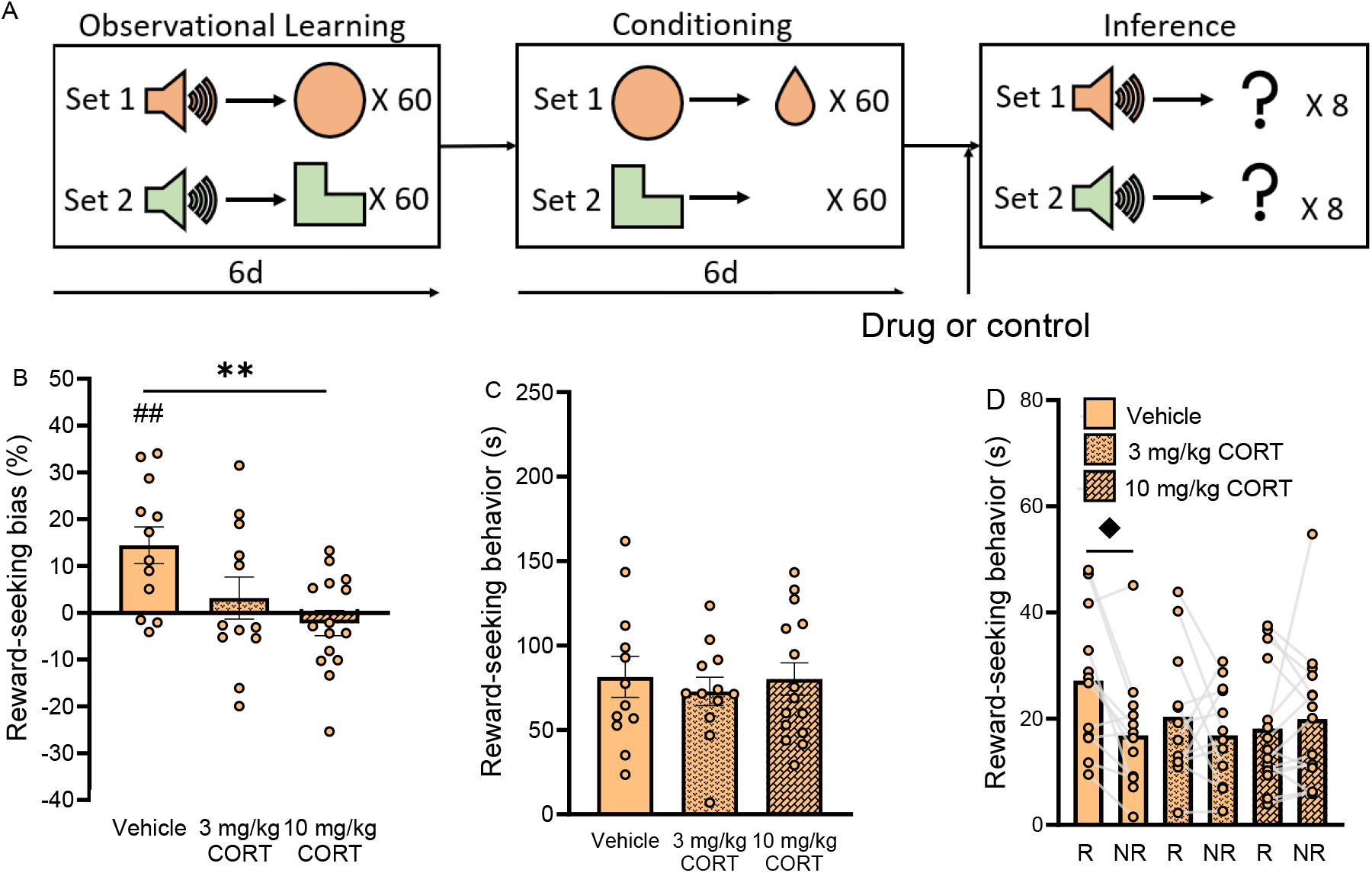
Systemic CORT administration impairs mnemonic inference, as reflected by a reduced reward-seeking bias on the memory inference test. **A.** Experimental timeline of the memory inference task. During the observational learning stage, mice were exposed to two different auditory-visual cue associations for 120 trials/day over 6 days. During the conditioning stage, they were trained to associate one of the visual cues with a sucrose reward for 120 trials/day over 6 days. During the memory inference test, 24 h after the last conditioning day, mice were re-exposed to the two auditory cues (8 trials per cue) and reward-seeking behavior was measured. Animals that infer correctly should preferentially seek reward to the auditory cue previously linked to the rewarded visual cue. CORT (3 or 10 mg/kg) or vehicle was administered intra-peritoneally 45 min prior to the memory inference test. **B**. CORT impaired the reward-seeking bias. **C**. CORT did not affect total reward-seeking behavior. **D**. The CORT-induced reduction in reward-seeking bias was caused by similar reward-seeking behavior to the rewarded (R) and non-rewarded (NR) set in animals treated with CORT. Vehicle: *n* = 12, 3 mg/kg CORT: *n* = 12, 10 mg/kg CORT: *n* = 15. Data represent mean ± SEM, circles represent individual data points. \*\**P* < 0.01 vs. vehicle, ^##^*P* < 0.01 vs. chance level, ^♦^*P* < 0.05 vs. non-rewarded set.

Vehicle-treated control mice showed successful inference, i.e. their reward-seeking bias was significantly above chance (one-sample *t*-test: *t*_*11*_ = 3.67, *P* = 0.004, Figure 1B). CORT significantly affected the reward-seeking bias (*F*_*2,36*_ = 5.42, *P* = 0.009) with the highest dose of CORT (10 mg/kg) significantly reducing the reward-seeking bias relative to vehicle controls (*t*_*23*_ = 5.10, *P* = 0.007), whereas 3 mg/kg CORT had no significant effect (*t*_*20*_ = 2.10, *P* = 0.129, Figure 1B). Total reward-seeking behavior did not differ between treatment groups (*F*_*2,36*_ = 0.20, *P* = 0.821; Figure 1C), nor did CORT treatment affect the reward-seeking bias when animals were re-exposed to the rewarded visual cues (*F*_*2,26*_ = 1.33, *P* = 0.282, Figure S1A), indicating that CORT treatment did not generally affect locomotor activity or incentive to seek reward.

Analysis of reward-seeking behavior to the two auditory cues separately revealed a significant CORT treatment × cue interaction effect (*F*_*2,36*_ = 3.48, *P* = 0.042), without main effects of CORT treatment (*F*_*2,36*_ = 0.40, *P* = 0.673) or cue (*F*_*2,36*_ = 4.09, *P* = 0.051; Figure 1D). Vehicle-treated control mice displayed higher reward-seeking behavior to the auditory cue associated with the rewarded set than to the non-rewarded set (*t*_11_ = 3.00, *P* = 0.014), whereas CORT-treated mice responded similarly to both auditory cues (3 mg/kg: *t*_11_ = 0.92, *P* = 0.999; 10 mg/kg: *t*_14_ = 0.60, *P* = 0.999). Tone discrimination remained intact (Figure S2). Thus, CORT-treated animals still inferred the auditory cue-outcome relationship but showed reduced inference accuracy, responding to both cues equally.

### Systemic yohimbine administration also impairs mnemonic inference

To examine whether noradrenergic activation also affected memory inference, yohimbine (1 or 3 mg/kg) or saline control was administered systemically 45 min before the memory inference test (Figure 1A).

Saline-treated control mice showed successful inference, i.e., the reward-seeking bias was significantly above chance (*t*_*16*_ = 4.57, *P* < 0.001). The highest dose of yohimbine (3 mg/kg) significantly reduced the reward-seeking bias relative to controls (one-way ANOVA: *F*_*2,39*_ = 8.32, *P* < 0.001; post-hoc: *t*_*31*_ = 3.58, *P* < 0.001; Figure 2A). Again, total reward-seeking behavior did not differ between treatment groups (*F*_*2,46*_ = 3.32, *P* = 0.047; Figure 2B), nor did yohimbine treatment affect the reward-seeking bias when animals were re-exposed to the rewarded visual cues (*F*_*2,30*_ = 0.61, *P* = 0.553; Figure S1B). Analysis of the response to the two auditory cues separately also revealed a significant yohimbine treatment × cue interaction effect (*F*_*2,39*_ = 4.84, *P* = 0.013), without main effects of yohimbine treatment (*F*_*2,38*_ = 2.15, *P* = 0.130) or cue (*F*_*2,38*_ = 0.62, *P* = 0.435). Saline-treated animals displayed higher reward-seeking behavior to the auditory cue associated with the rewarded set than to that of the non-rewarded set (*t*_*16*_ = 2.76, *P* = 0.031), whereas yohimbine-treated animals did not (1 mg/kg: *t*_8_ = 0.37, *P* = 0.999; 3 mg/kg: *t*_15_ = 1.65, *P* = 0.999, Figure 2C). Thus, similar to CORT treatment, yohimbine did not prevent animals from inferring the novel association *per se*. Rather, yohimbine also caused animals to seek reward to both cues equally, revealing a deficit in the accuracy of the inference.

**Figure 2.**
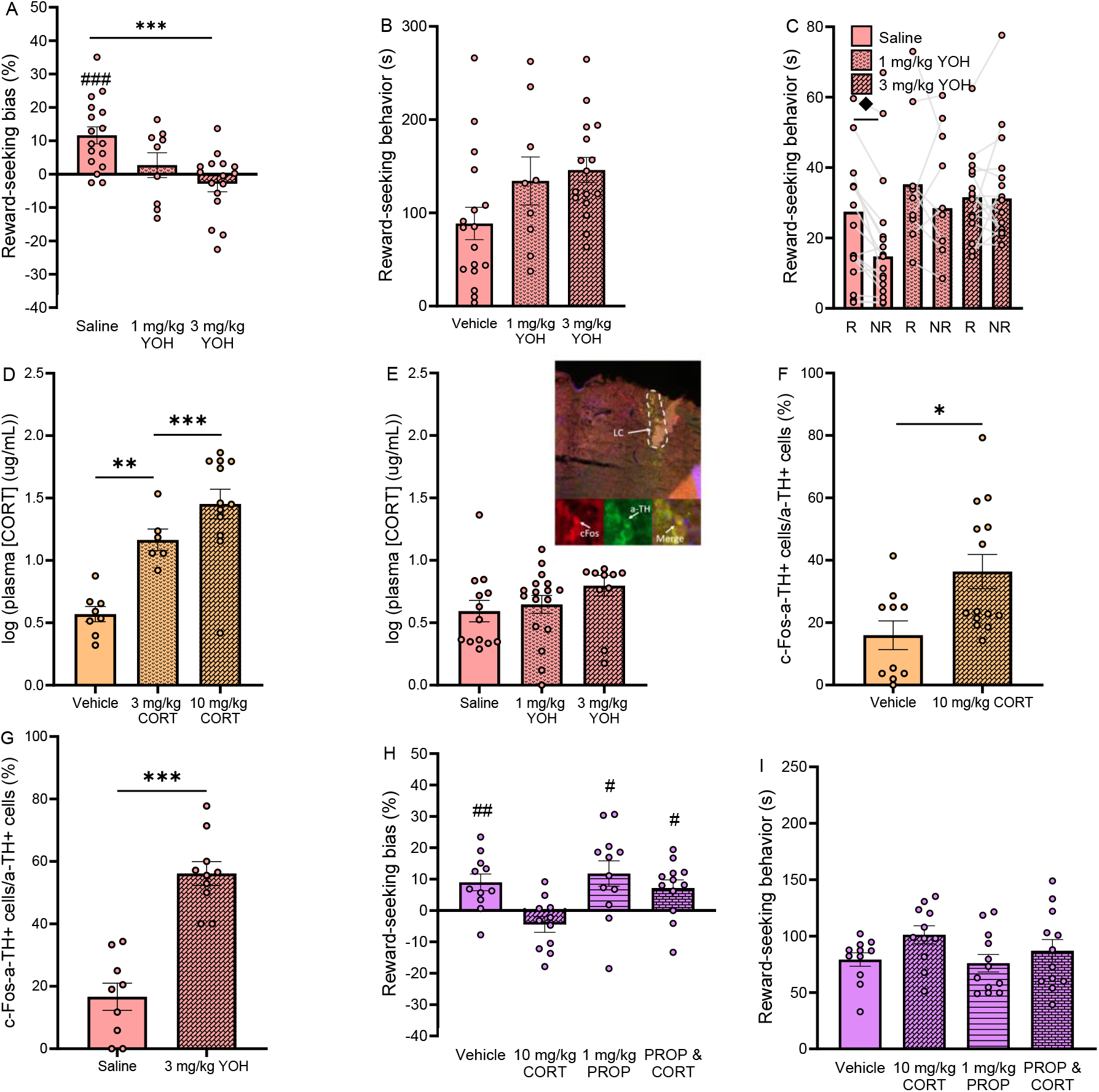
CORT administration affects noradrenergic signaling and may disrupt mnemonic inference. **A.** Yohimbine impaired the reward-seeking bias. **B**. Yohimbine did not affect total reward-seeking behavior. **C**. The yohimbine-induced impairment in reward-seeking bias was caused by similar reward-seeking behavior to the rewarded (R) and non-rewarded (NR) set. Saline: *n* = 17, 1 mg/kg yohimbine: *n* = 9, 3 mg/kg yohimbine: *n* = 16. **D**. CORT administration increased plasma CORT levels (3 mg/kg: *t*_*13*_ = 0.17, *P* = 0.006; 10 mg/kg: *t*_*19*_ = 6.16, *P* < 0.001, relative to vehicle). Vehicle: *n* = 8, 3 mg/kg CORT: *n* = 6, 10 mg/kg CORT: *n* = 12. **E**. Yohimbine did not increase plasma CORT levels. Saline: *n* = 14, 1 mg/kg YOH: *n* = 12, 3 mg/kg YOH: *n* = 14. **F**. Yohimbine administration increased c-Fos-expression in noradrenergic neurons (marked by tyrosine hydroxylase expression) of the locus coeruleus (*t*_*17*_ = 6.23, *P* < 0.001). Saline: *n* = 9, 3 mg/kg YOH: *n* = 10. **G**. CORT administration also increased c-Fos-expression in noradrenergic neurons (marked by tyrosine hydroxylase expression) of the locus coeruleus (*t*_*22*_ = 2.70, *P* = 0.013). Vehicle: *n* = 10, 10 mg/kg CORT: *n* = 14. **H**. Propranolol reduced the CORT-induced impairment in reward-seeking bias; CORT administration alone resulted in a reward-seeking bias not significantly different from 0. **I**. Propranolol or CORT + propranolol treatment had no effect on total reward-seeking behavior (CORT: *F*_*3,42*_ = 5.77, *P* = 0.221, propranolol: *F*_*3,42*_ = 2.15, *P* = 0.151, CORT × propranolol interaction: *F*_*3,42*_ = 0.06, *P* = 0.814). Vehicle: *n* = 11, 10 mg/kg CORT: *n* = 11, 1 mg/kg PROP: *n* = 12, CORT + PROP: *n* = 12. Data represent mean ± SEM, circles represent individual data points. \**P* < 0.05, \*\**P* < 0.01, *** *P* < 0.001 vs. control, ^#^*P* < 0.05, ^##^*P* < 0.01, ^###^*P* < 0.001 vs. chance level, ^♦^*P* < 0.05 vs. non-rewarded set.

### Noradrenergic signaling is affected by CORT administration and may disrupt mnemonic inference

As we found that CORT and yohimbine induced highly comparable effects on memory inference, we next investigated whether these two stress hormone systems interact in influencing memory inference. We first examined whether CORT or yohimbine administration increased plasma CORT levels in blood collected 30 min after the memory inference test. CORT increased plasma CORT levels (*F*_*2,23*_ = 19.05, *P* < 0.001), whereas yohimbine administration did not (*F*_*2,37*_ = 2.27, *P* = 0.118; Figure 2D-E). Interestingly, both CORT and yohimbine treatment increased activity of noradrenergic neurons (marked by tyrosine hydroxylase expression) of the locus coeruleus 30 min after the memory inference test (CORT: *t*_*22*_ = 2.70, *P* = 0.013; yohimbine: *t*_*17*_ = 6.23, *P* < 0.001; Figure 2F-G, Figure S3A-B). These findings indicate that systemic CORT administration enhanced activity of the noradrenergic system.

Next, to examine if the CORT effect on memory inference is dependent on noradrenergic activity, we systemically administered the β-adrenoceptor antagonist propranolol (1 mg/kg) together with CORT (10 mg/kg) 45 min before testing. Importantly, while CORT alone impaired the reward-seeking bias (one-sample *t*-test: *t*_10_ = 1.76, *P* = 0.113), CORT-propranolol co-administration (*t*_11_ = 2.72, *P* = 0.022) resulted in a reward-seeking bias significantly above chance, same as in vehicle (*t*_10_ = 3.36, *P* = 0.007) and propranolol only groups (*t*_11_ = 2.92, *P* = 0.013). While this pattern suggests that propranolol administration may prevent the CORT-induced inference deficit, this finding has to be interpreted with caution as the CORT × propranolol treatment interaction did not reach significance (*F*_*3,42*_ = 0.70, *P* = 0.408, Figure 2H). Total reward-seeking behavior was not affected (all *P*’s > 0.151, Figure 2I).

### CORT reduces pyramidal cell activity and increases GABAergic cell activity in area CA3

We next examined *c-Fos* mRNA expression 30 min after the memory inference test in the dentate gyrus (DG), and the stratum radiatum and pyramidal cell layers of CA1 (CA1sr, CA1py) and CA3 (CA3sr, CA3py), using FISH. In CORT-treated (10 mg/kg) mice, we found a significant decrease in the total number of *c-Fos*-expressing cells in CA3py (*t*_*16*_ = 3.04, *P* = 0.024, Figure 3B), but not in the DG (*t*_*16*_ = 0.87, *P* = 0.395) or CA1py (*t*_*16*_ = 0.21, *P* = 0.834). Further, CORT treatment increased *c-Fos* activity within GABAergic neurons of the CA3sr (*t*_*16*_ = 3.95, *P* = 0.003, Figure 3C), but not in the DG (*t*_*16*_ = 0.61, *P* = 0.554) or CA1sr (*t*_*16*_ = 0.40, *P* = 0.693). Also, CORT treatment did not significantly affect *c-Fos* within glutamatergic neurons in any hippocampal subregion (*P*’s > 0.180, Figure 3D). Total numbers of GABAergic or glutamatergic cells remained unchanged (*P*’s > 0.101; Figure S4).

**Figure 3.**
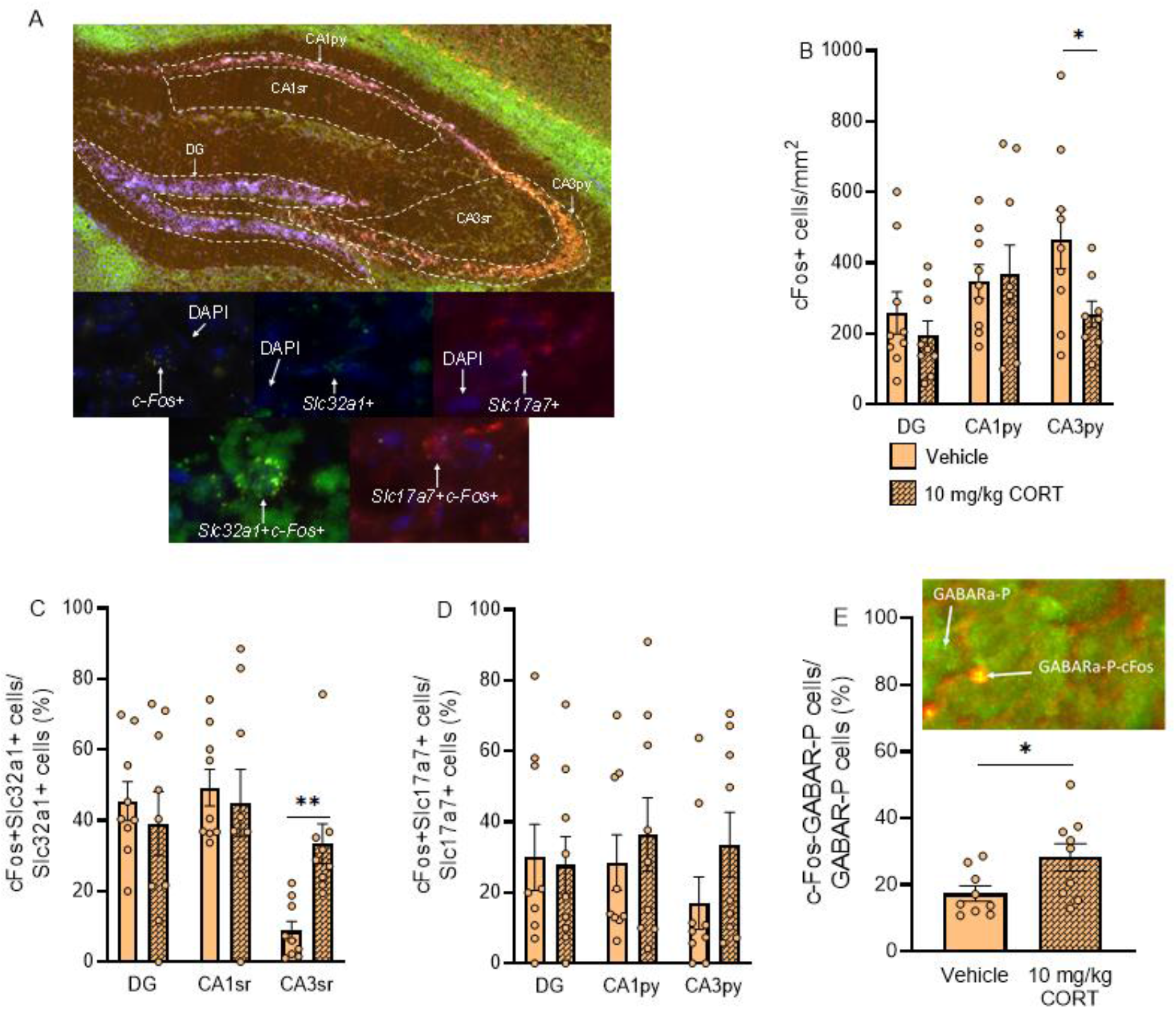
CORT reduces pyramidal cell activity and increases GABAergic cell activity in area CA3. **A.** Images from an exemplary slice showing the hippocampal subregions and quality of FISH staining (DAPI+, *c-Fos*+, *Slc32a1*+, and *Slc17a7*+ mRNA). **B**. CORT reduced the number of *c-Fos*-expressing cells in area CA3. **C**. CORT increased the number of neurons showing *c-Fos*-expression in GABAergic cells (*Slc32a1*- expressing) in area CA3. **D**. CORT did not affect the number of neurons showing *c-Fos* expression in glutamatergic cells (*Slc17a7*-expressing) in the hippocampus. Vehicle: *n* = 9, 10 mg/kg CORT: *n* = 9. **E**. CORT increased phosphorylated GABA_A_ receptor levels in c-Fos-expressing cells. Vehicle: *n* = 9, 10 mg/kg CORT: *n* = 9. Data represent mean ± SEM, circles represent individual data points. \**P* < 0.05, \*\**P* < 0.01 vs. vehicle.

To investigate whether enhanced activity of GABAergic neurons underlies reduced CA3py neuronal activity, we examined GABA_A_ receptor activity in CA3 pyramidal cells^32,33^. CORT increased GABA_A_ receptor phosphorylation in c-Fos-expressing CA3py cells (*t*_*16*_ = 2.31, *P* = 0.034, Figure 3E), indicating enhanced inhibitory signaling.

Notably, these CORT-induced changes in CA3 activity were absent when animals were re-exposed to the visual cues alone (*P*’s > 0.101, Figure S1C-D), or when animals were tested on a 2-tone recall paradigm, where animals were trained to associate auditory cues with reward, and subsequently re-exposed to the auditory cues (Figure S2). These findings indicate that CORT does not affect CA3 neuronal activity when animals simply recall learned information. Rather, a CORT-induced reduction in CA3 activity seems to underlie impaired accuracy in the inference test. Indeed, CA3 activity was not altered by CORT-treatment when differentiation between rewarded and non-rewarded set was not needed, i.e., both visual cues were rewarded during conditioning, (Figure S5). Moreover, similar CA3 patterns emerged when animals were grouped by the reward-seeking bias (i.e., good performers >5%; bad performers <5%) rather than treatment, using a separate immunohistochemistry protocol with GABAergic marker glutamate decarboxylase 67 (GAD67; Figure S6). Together, these findings indicate that this CORT-induced enhanced GABAergic signaling in CA3 is a potential mechanism of impaired memory inference performance.

### Yohimbine reduces pyramidal cell activity and increases GABAergic cell activity in area CA3

We next examined whether yohimbine (3 mg/kg) similarly influenced hippocampal neuronal activity. In yohimbine-treated mice, we found a significant decrease in total *c-Fos*-expression in the CA3py (*t*_*16*_ = 4.04, *P* = 0.003; Figure 4A), but not in the CA1py (*t*_*16*_ = 0.28, *P* = 0.785) or DG (*t*_*16*_ = 0.14, *P* = 0.895). Further, yohimbine treatment increased *c-Fos* activity within GABAergic neurons of the CA3sr (*t*_*16*_ = 3.65, *P* = 0.006, Figure 4B), whereas this effect was absent in CA1sr (*t*_*16*_ = 0.32, *P* = 0.751) or DG (*t*_*16*_ = 0.19, *P* = 0.853). Again, no differences were found in *c-Fos* activity within glutamatergic neurons in any hippocampal subregion (*P*’s > 0.233, Figure 4C). Total numbers of GABAergic or glutamatergic cells remained unchanged (*P*’s > 0.101; Figure S4). Thus, similar to CORT, yohimbine also induces enhanced GABAergic signaling in CA3, a potential mechanism of the deficit in memory inference accuracy.

**Figure 4.**
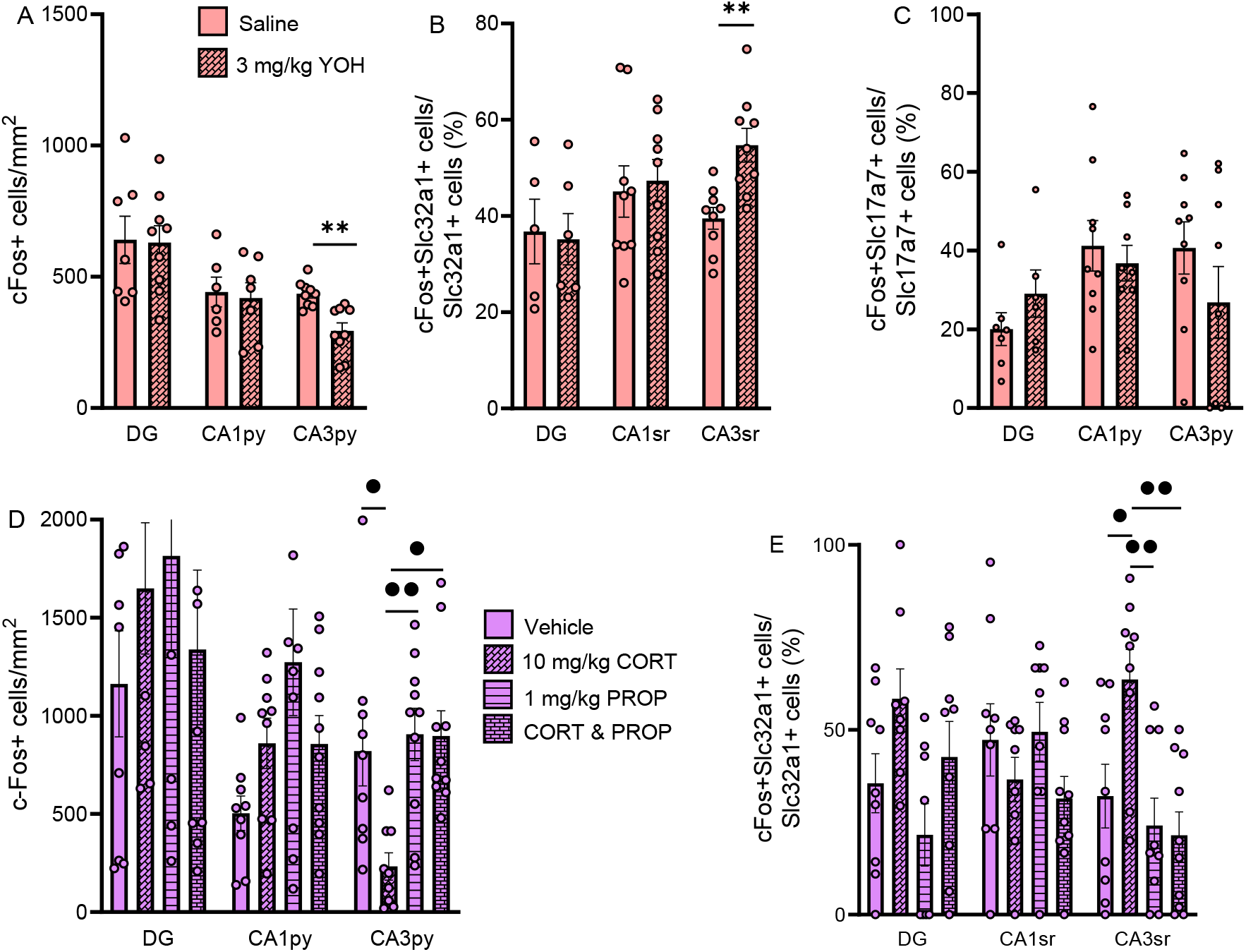
Propranolol prevents the CORT-induced effects on neuronal activity in area CA3. **A.** Yohimbine reduced *c-Fos*-expression in area CA3. **B**. Yohimbine increased *c-Fos*-expression in GABAergic cells in area CA3. **C**. Yohimbine did not affect *c-Fos*-expression in glutamatergic cells in the hippocampus. Saline: *n* = 9, 3 mg/kg yohimbine: *n* = 10. **D**. Propranolol co-administration prevented the CORT-induced reduction in CA3 activity. *c-Fos*-expression was not altered in area CA1 (CORT: *F*_*3,33*_ = 0.02, *P* = 0.888; propranolol: *F*_*3,33*_ = 4.28, *P* = 0.046; CORT × propranolol interaction: *F*_*3,33*_ = 04.36, *P* = 0.045; post-hoc test: *P* > 0.096), or in the DG (CORT: *F*_*3,33*_ = 0.0017, *P* = 0.990; propranolol: *F*_*3,33*_ = 0.20, *P* = 0.660; CORT × propranolol interaction: *F*_*3,33*_ = 1.57, *P* = 0.220). Notably, *c-Fos-*expression in area CA3 was not affected by CORT-propranolol co-administration relative to vehicle (*t*_*18*_ = 0.41, *P* = 0.999) or propranolol-administration (*t*_*18*_ = 0.02, *P* = 0.999). **E**. CORT and propranolol co-administration also prevented the CORT-induced increase *c-Fos*-expression in GABAergic cells in CA3. *c-Fos*-expression in GABAergic cells was not altered in area CA1 (CORT: *F*_*3,33*_ = 3.66, *P* = 0.064; propranolol: *F*_*3,33*_ = 0.04, *P* = 0.845; CORT × propranolol interaction: *F*_*3,33*_ = 0.24, *P* = 0.628), or in the DG (CORT: *F*_*3,33*_ = 6.53, *P* = 0.016 (no post-hoc effect); propranolol: *F*_*3,33*_ = 2.98, *P* = 0.095; CORT × propranolol interaction: *F*_*3,33*_ = 0.01, *P* = 0.919). Notably, *c-Fos-*expression in GABAergic cells in area CA3 was not affected by CORT-propranolol co-administration relative to vehicle (*t*_*18*_ = 1.00, *P* = 0.999) or propranolol-administration (*t*_*18*_ = 0.25, *P* = 0.999). Vehicle: *n* = 9, 10 mg/kg CORT: *n* = 9, 1 mg/kg PROP: *n* = 9, CORT + PROP: *n* = 10. Data represent mean ± SEM, circles represent individual data points. \**P* < 0.05 vs. saline, ^•^*P* < 0.05, ^••^*P* < 0.01 vs. 10 mg/kg CORT.

### Propranolol prevents the CORT-induced increase in GABAergic cell activity in area CA3

As both stress hormones induced comparable effects on hippocampal activity, we examined whether propranolol (1 mg/kg), administered 45 min prior to the inference test, prevents the effects of CORT administration on neuronal activity. In CA3, two-way ANOVA revealed significant effects of CORT (*F*_*3,33*_ = 4.57, *P* = 0.040), propranolol (*F*_*3,33*_ = 7.31, *P* = 0.011), and a significant CORT × propranolol interaction (*F*_*3,33*_ = 4.72, *P* = 0.037; Figure 4D). CORT reduced total *c-Fos*-expression relative to vehicle-treated animals (*t*_*17*_ = 3.10, *P* = 0.030), propranolol-treated animals (*t*_*17*_ = 3.38, *P* = 0.008), and CORT-propranolol co-administered animals (*t*_*18*_ = 3.38, *P* = 0.011; Figure 4D).

Similarly, two-way ANOVA for GABAergic activity in CA3sr indicated a significant CORT x propranolol interaction effect (*F*_*3,33*_ = 5.06, *P* = 0.031; main effect propranolol: *F*_*3,33*_ = 10.89, *P* = 0.002; main effects CORT: *F*_*3,33*_ = 3.62, *P* = 0.066; Figure 4E). CORT increased relative GABAergic activity relative to vehicle-treatment (*t*_*17*_ = 2.90, *P* = 0.040), propranolol treatment (*t*_*17*_ = 3.63, *P* = 0.006), and CORT-propranolol co-administration (*t*_*18*_ = 3.98, *P* = 0.002; Figure 4E), indicating that propranolol prevented CORT-induced increases in *c-Fos*-expression within GABAergic neurons in area CA3sr.

### GABA_A_ receptor antagonist bicuculline prevents the CORT-induced impairment in reward-seeking bias

We found that CORT impaired inference accuracy and increased GABAergic signaling in hippocampal area CA3. To investigate whether GABAergic signaling within the hippocampus is necessary to mediate the CORT effect on memory accuracy, we administered CORT (3 ng) together with the GABA_A_ receptor antagonist bicuculline methiodide (0.5 ng) into the hippocampus 45 min before inference testing (Figure 5A). Two-way ANOVA revealed a significant main effect of bicuculline (*F*_*3,57*_ = 9.76, *P* = 0.003) and a CORT × bicuculline interaction (*F*_*3,57*_ = 8.46, *P* = 0.005), but no main effect of CORT (*F*_*3,57*_ = 1.33, *P* = 0.255; Figure 5B). CORT impaired the reward-seeking bias relative to vehicle (*t*_*31*_ = 2.99, *P* = 0.015), whereas bicuculline blocked this impairment, with CORT + bicuculline-treated animals differing significantly from CORT-treated animals (*t*_*29*_ = 4.27, *P* < 0.001), but not from vehicle- (*t*_*31*_ = 1.34, *P* = 0.999), or bicuculline-treated controls (*t*_*23*_ = 1.20, *P* = 0.999; Figure 5B). Bicuculline alone did not affect the reward-seeking bias (*t*_*28*_ = 0.15, *P* = 0.999).

**Figure 5.**
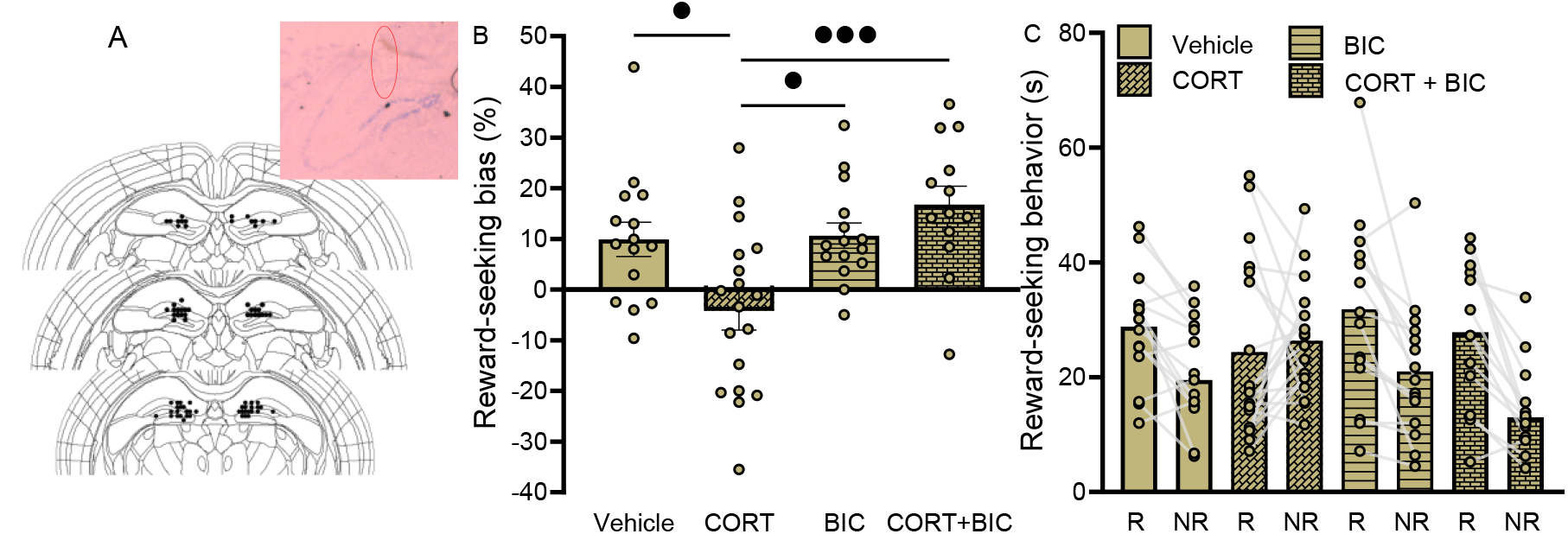
GABA_A_ receptor antagonist bicuculline prevents the CORT-induced impairment in reward-seeking bias. **A.** Histological image of an exemplary brain showing the position of the needles. The coordinates of the needle tips are also shown. **B**. Bicuculline (BIC: 0.5 ng) prevented the CORT (3 ng) induced impairment in reward-seeking bias. **C**. Bicuculline or CORT treatment had no effect on total reward-seeking behavior. Vehicle: *n* = 15, 3 ng CORT: *n* = 18, 0.5 ng BIC: *n* = 15, CORT & BIC: *n* = 13. Data represent mean ± SEM, circles represent individual data points. ^•^*P* < 0.05, ^•••^*P* < 0.001 vs. CORT.

Total reward-seeking behavior was unaffected by CORT (*F*_*3,57*_ = 0.38, *P* = 0.542), bicuculline (*F*_*3,57*_ = 1.72, *P* = 0.196) or their interaction (*F*_*3,57*_ = 2.91, *P* = 0.093). Further, CORT treatment did not alter responding to rewarded visual cues (*F*_*2,27*_ = 0.96, *P* = 0.650; Figure S7), or plasma CORT levels (*F*_*2,36*_ = 0.12, *P* = 0.887, Figure S7). Thus, these findings indicate that the CORT-induced increase in GABAergic activity in the hippocampus plays a critical role in mediating the CORT effect on impairing memory inference.

## Discussion

The ability to integrate discrete events that share common elements allows the formation of knowledge structures that facilitate prediction in complex environments and is thus fundamental for adaptive behavior. Recent work in humans has shown that stress can impair memory inference^10^, but the underlying mechanism remained unknown. Here, we demonstrate that CORT reduced memory inference accuracy through a noradrenergic-dependent mechanism involving increased GABAergic inhibition and reduced CA3 activity. Blocking β-adrenoceptors or GABA_A_ receptors prevented this CORT-induced deficit, identifying a circuit through which stress hormones impair mnemonic inference accuracy.

CORT selectively impairs mnemonic inference accuracy. Mice first learned two sets of auditory-visual associations, after which one visual cue was rewarded. At test, vehicle-treated control mice selectively sought reward to the auditory cue associated with the rewarded visual cue. CORT-treated mice sought reward in response to both auditory cues, indicating reduced inference accuracy. As stress and CORT can impair memory retrieval^2,34,35^, one potential explanation is that CORT interfered with memory retrieval. However, our data suggest that retrieval deficits alone cannot account for accuracy impairments, as CORT-treated mice still responded to auditory stimuli and correctly sought rewarded to previously rewarded visual cues. These data agree with human findings, indicating that stress-related inference impairments cannot be fully explained by deficits in retrieval alone^11^.

Importantly, inference can arise during encoding of overlapping content (integrative encoding)^8,36^, or during retrieval^37,38^. In humans, memory integration is associated with increased hippocampus-prefrontal cortex activity^8,9,37,38^, allowing online reactivation and recombination of memories.

Consistent with this view, successful inference has been shown to depend on reactivation of the associated element during learning of overlapping content^11,14,38^. Recent human studies show that stress before encoding disrupts integrative encoding and impairs later inference^11^, whereas our findings show that stress hormone administration (or stress exposure in recent human work^10^) before retrieval also impairs inference. Thus, while stress before encoding impairs integrative encoding, resulting in later inference deficits^11^, stress before inference testing can result in inference deficits as well.

The hippocampus contributes to both encoding and recall processes^39–41^, and is thought to play a key role in mnemonic inference^8,9^. In line with this idea, our findings demonstrate that CORT reduced CA3py activity, an effect that was absent in non-inference conditions. Given the established role of CA3 in pattern completion in mnemonic inference^14,42^, this finding points to a potential mechanism by which stress hormones impair inference accuracy. Previous studies emphasize a role of the CA1 in inference^8,38^, but these studies primarily assessed hippocampal functioning for memory inference *per se*. In contrast, our findings suggest that area CA3 contributes to the accuracy of inference. Consistent with this interpretation, when groups were split on inference performance rather than treatment, the reduction in CA3 activity persisted, indicating that diminished CA3 activity is specifically associated with reduced inference accuracy rather than an impaired ability to infer.

Reduced CA3 pyramidal neuronal activity was coupled to an increase in *c-Fos* expression in GABAergic cells in the CA3sr, suggesting that GABAergic activity in CA3sr might reduce CA3 pyramidal cell activity. CA3sr axons reach into the pyramidal layer^43^, and activation of CA3sr interneurons has been shown to suppress cellular activity in the pyramidal layer^44^. Further, at the synaptic level, we observed increased phosphorylation of the GABA_A_ receptor in CA3py. Although indirect, increased phosphorylation of the β3-subunit of the GABA_A_ receptor is consistent with enhanced GABAergic signaling^32,33^. Moreover, administration of the GABA_A_ receptor antagonist bicuculline methiodide prevented the CORT-induced impairment of memory inference accuracy. Thus, together these findings support a model in which increased GABAergic inhibition within area CA3 suppresses pyramidal neuronal activity, thereby impairing inference accuracy.

Yohimbine produced inference deficits similar to CORT, consistent with a shared noradrenergic mechanism. Interneurons within the hippocampus express α_1_- and β-adrenoceptors, as well as glucocorticoid receptors^26,27^. Activation of these receptors by norepinephrine or CORT has been shown to enhance GABAergic signaling within the hippocampus^26,27^, suggesting a possible mechanism for the observed increase in inhibition. Importantly, CORT and norepinephrine interact in regulating memory, with CORT facilitating noradrenergic effects in rodents^22,23,45^ and humans^46–48^. Similar interaction effects have also been reported in memory flexibility studies^49–52^. In line with these findings, we show that blockade of β-adrenoceptors with propranolol prevents the CORT-induced impairment in inference accuracy, indicating that this deficit in memory flexibility is dependent on noradrenergic signaling. These findings indicate that CORT impairs inference partly through noradrenergic enhancement of GABAergic inhibition in CA3.

Thus, we demonstrate that CORT (and noradrenergic arousal) can interfere with mnemonic inference and identify a key mechanism underlying this effect. CORT, through noradrenergic signaling, disrupts CA3-dependent processes by increased GABAergic signaling within this region. Accordingly, blocking either GABAergic or noradrenergic activity diminished the CORT-induced inference deficit. Importantly, our findings significantly extend recent human work on the impact of stress on mnemonic inference^10,11^ by providing insights into the underlying neuroendocrine mechanisms. Our findings identify memory inference as a key cognitive process vulnerable to major stress mediators. By disrupting the integration of overlapping memories, stress may distort how relationships between events are inferred, limiting behavioral flexibility and adaptation to complex environments. Given that similar deficits are observed in mental disorders such as post-traumatic stress disorder^47^ and anxiety^53,54^, impaired memory inference may represent an important cognitive mechanism linking stress to psychopathology. Our findings point to potential targets (i.e. noradrenergic and GABAergic signaling) for treating such stress-related inference deficits.

## Data Availability Statement

All data will be made available upon request.

## Notes

### Competing Interest Statement

The authors have declared no competing interest.

### Funding

- Deutsche Forschungsgemeinschaft: 449640848

