## Supplementary Methods for "Glucocorticoids impair memory inference through noradrenergic disruption of GABAergic regulation in hippocampal CA3"

### Memory inference task

Prior to training, mice were handled for 4 min per day for 3 days to become accustomed to the experimenter. Training and testing were performed in a dark, sound-proof triangular box (depth: 170 mm; height: 230 mm; width reward side: 46 mm; width screen side: 238 mm, Campden, Bussey-Saksida Touch Screen system model 80614-20). Boxes were equipped with a screen, speaker and reward module. Boxes were cleaned with 40% ethanol between animals.

The memory inference task consisted of three stages: observational learning, conditioning, and memory inference testing (Figure 1A; Barron et al., 2020). In the first stage (observational learning), animals were exposed to two sets of auditory cue-visual cue associations: a 2-kHz tone paired to an orange circle (set 1), and a 500-Hz tone paired to a green L-shape (set 2). In a single trial, animals were exposed to the auditory cue for 10 s, followed by an 8-s exposure to the visual cue and an intertrial interval (ITI) of 60 s. Auditory cues were presented via a speaker and visual cues were presented on a screen. This stage consisted of 120 trials (60 trials of set 1, 60 trials of set 2, in a randomized order) per day for 6 days. Each daily training session lasted approximately 2.5 h.

In the second stage (conditioning), animals were exposed again to the two visual cues, but now one of these cues (i.e. either set 1 or set 2) was consistently rewarded by a 15% sucrose droplet (100 µL) from a dispenser in the outcome area. Animals were exposed to the visual cue for 8 s, followed by delivery of reward or nothing and an ITI of 60 s. This stage consisted of 120 trials per day for 6 days. On the first day, sets were presented in blocks of 6 trials (20 blocks in total); on the other days, sets were presented in a random order (60 trials of set 1 and 60 trials of set 2). Each daily training session lasted approximately 2.5 h.

During the final stage (memory inference test), animals were exposed to both auditory cues in isolation for 10 s, and reward-seeking behavior was quantified as the time spent in the outcome area during the 20 s following cue termination. The test consisted of 16 trials (8 per set) with a 30 s ITI. Increased reward-seeking behavior for one of the two sets (which was rewarded during the conditioning) indicated that animals had inferred the novel association between auditory cue and outcome. After memory inference testing, animals were placed back in their home cage.

Animals' behavior was recorded with a video camera mounted above the experimental apparatus. Videos were analyzed offline by a trained observer blind to the treatment condition, and reward-seeking behavior was scored. Reward-seeking behavior was defined as the time spent near the reward area (nose towards the reward module at <2 cm) in the 20-s period after termination of the cue. A reward-seeking bias was computed by subtracting the reward-seeking behavior of the non-rewarded set from the reward-seeking behavior of the rewarded set and dividing this by the total reward-seeking behavior (i.e. [reward-seeking behavior rewarded – reward-seeking behavior non-rewarded]/[reward-seeking behavior rewarded + reward-seeking behavior non-rewarded + reward-seeking behavior ITI] × 100 %). Animals were excluded (in total 12% of animals were excluded) from analyses when they did not display a reward-seeking bias higher than 0 during the last day of conditioning (i.e. did not acquire the visual cue-outcome association) or showed abnormal total reward-seeking behavior during the test (>2 standard deviations from the mean).

As drug was administered shortly before the memory inference test, it could also non-selectively influence performance. Therefore, we implemented several control conditions.

*Motivation to seek reward – visual cues and reward only*

First, to examine whether drug affected general motivation to seek reward, mice were trained on the task as described before, but during the test were re-exposed to the two visual cues (8 trials of set 1, 8 trials of set 2) followed by reward or nothing. During this final stage, animals were exposed to each visual cue in isolation for 10 s, followed by a sucrose reward, and reward-seeking behavior was quantified as the time spent in the outcome area during the 20 s following cue termination. This stage consisted of 16 trials (8 trials per set) with an ITI of 30 s. After testing animals were placed back in their home cage.

*Recall – auditory cue-reward associations*

To determine whether the drug manipulation had affected recall of the learned cue-outcome associations, a memory recall task was performed. This task consisted of two stages: conditioning, and recall testing (Figure S2A). Two days prior to the start of the first stage, animals were food restricted to 90% of their free-feeding body weight. In the first stage (conditioning), animals were exposed to two auditory cues (set 1: 2 kHz, set 2: 500 Hz), and one of these cues (i.e. either set 1 or set 2) was consistently rewarded by a 15% sucrose droplet (100 μL) from a dispenser in the outcome area. Animals were exposed to the auditory cue for 8 s, followed by delivery of reward or nothing and an ITI of 60 s for 120 trials per day for 6 days. Sets were presented in random order (60 trials of set 1 and 60 trials of set 2). During the final stage (recall testing), animals were exposed to each auditory cue in isolation for 10 s, and reward-seeking behavior was quantified as the time spent in the outcome area during the 20 s following cue termination. This stage consisted of 16 trials (8 trials per set) with an ITI of 30 s. After recall testing animals were placed back in their home cage.

*Discrimination – rewarded-and-novel auditory cue*

Lastly, to determine whether drug administration might have affected the animals' ability to discriminate between the different auditory cues, another control experiment was performed. On days 3 and 5 of the conditioning stage in the memory recall task (as described before), animals were exposed to the rewarded cue (either set 1 or set 2) and a novel auditory cue of 3 kHz (Figure S2A). Vehicle or drug was administered in a counterbalanced order (i.e. if an animal received drug on day 3, it received vehicle on day 5 and vice versa). Animals were exposed to each auditory cue for 8 s, followed by reward delivery or nothing. Cues were presented in a random order for 10 trials (5 rewarded cue, 5 novel cue) with an ITI of 60 s. After 10 trials, training proceeded as described above. Reward-seeking behavior was quantified as the time spent in the outcome area during the 20 s following cue termination.

As we found a stress hormone-induced reduction in memory inference accuracy, we performed two more experiments to examine the extent to which stress hormones reduced inference accuracy and whether this was dependent on the hippocampus.

*Extension to novel auditory cues – novel cue experiment*

Animals were trained on the memory inference task as described before (i.e. observational learning and conditioning stages; Figure 1A). Twenty-four hours after the last training session, animals were administered vehicle or CORT and, 45 min later, re-exposed to the auditory cue of the rewarded set (i.e. either 500 Hz or 2 kHz) and a novel auditory cue of 3 kHz. Animals were exposed to both auditory cues in isolation for 10 s, and reward-seeking behavior was quantified as the time spent in the outcome area during the 20 s following cue termination. The test consisted of 16 trials (8 per set) with a 30 s ITI. After testing, animals were placed back in their home cage.

*Accuracy – both visual cues rewarded*

Next, to examine whether the stress-induced alteration in hippocampal activity was due to animals having to choose between the two sets, we trained animals on the memory inference task as described before, but rewarded both visual cues with a 15% sucrose reward during the conditioning stage. This removed the need for animals to choose during the inference test. Forty-five minutes before the inference test, CORT or vehicle was systemically administered, and animals were re-exposed both auditory cues in isolation for 10 s. Reward-seeking behavior was quantified as the time spent in the outcome area during the 20 s following cue termination. The test consisted of 16 trials (8 per set) with a 30 s ITI. Increased reward-seeking behavior for both sets (as both visual cues were rewarded during conditioning) indicated that animals had inferred the novel association between auditory cue and outcome. After testing, animals were placed back in their home cage.

**Fluorescent in-situ hybridization**

Thirty minutes after the memory inference test, animals were killed by cervical dislocation, the brains rapidly dissected, and frozen in isopentane to obtain fresh tissue for RNA measurements. Twenty-micrometer thick coronal sections of the brains were made on a cryostat (Leica), and mounted directly on SuperFrost Plus slides. Sections were left to dry at -20 °C for 2 h, and stored at -70 °C for later use.

Every fourth section of the dorsal hippocampus (AP: -1.46 mm to -2.30 mm, relative to Bregma) was selected for later staining (four sections per animal). A fluorescent in-situ hybridization (FISH) protocol was performed using the standard protocol of Advanced Cell Diagnostics (ACD; RNAscope™ Multiplex Fluorescent V2 Assay kit, cat: 323100). To examine *c-Fos* expression in both glutamatergic and GABAergic neurons, probes against *c-Fos* (Mm-Fos-Mus Musculus FBJ osteosarcoma oncogene (*Fos*), mRNA; ACD #316921), *Slc17a7* (Mm-Slc17a7-O2-C2 - Mus musculus solute carrier family 17 (sodium-dependent inorganic phosphate cotransporter) member 7 (*Slc17a7*) mRNA; ACD #501101-C2), and *Slc32a1* (Mm-Slc32a1-C3 – Mus Musculus solute carrier family 32 (GABA vesicular transporter) member 1 (*Slc32a1*) mRNA; ACD #31991-C3) mRNA were used. Sections were first post-fixed in 4% paraformaldehyde (PFA, pH 7.1 – 7.2) for 75 min at 4 °C, and dehydrated in 50, 70 and 100% ethanol in MilliQ for 5 min each at room temperature. Sections were then stored in fresh 100% ethanol at -20 °C overnight. The next day, sections were incubated with 5% hydrogen peroxide for 10 min at room temperature, washed in MilliQ twice for 1 min and baked at 37 °C in an oven for 30 min to remove the last hydrogen peroxide. As proteins can degrade mRNA, sections were incubated in protease for 15 min at room temperature. Next, sections were incubated with the probe mix (*Slc17a7* and *Slc32a1* probes were combined into a single probe mixture and diluted 1:50 with *c-Fos* probe diluent) for 2 h at 40 °C in a humidified chamber. Samples were then washed in wash buffer (0.1 M saline-sodium citrate buffer, 0.03% lithium dodecyl sulfate in 0.1 M phosphate-buffered saline (PBS)) twice for 2 min each. Sections were stored overnight in 5X saline-sodium citrate buffer at room temperature. The next day, sections were washed in wash buffer for 2 min, and signals were amplified using the provided amplification solutions (AMPs) for 30 min at 37°C for each channel (AMP-1, AMP-2), sections were incubated in AMP-3 for 15 min at 37 °C. Between AMP incubations, sections were washed twice in wash buffer for 2 min each. Afterwards, the fluorophore binding site was opened by incubating with horseradish peroxidase (HRP) reagents (HRP-C1, HRP-C2, HRP-C3) for 15 min at 40 °C and the tyramide signal amplifier (TSA) fluorophores were added: TSA 570 for channel 1 (*c-Fos*, ACD), TSA 650 for channel 2 (*Slc17a7*, ACD) and TSA 520 for channel 3 (*Slc32a1*, ACD), incubated for 30 min at 40 °C. The fluorophore site was closed using the provided HRP blocker for 15 min at 37 °C, followed by the opening of the next fluorophore site. Between each step, sections were washed in wash buffer 3 times for 2 min each. All steps were performed in a humidified chamber at 40 °C in the dark. Finally, sections were incubated in 4',6-diamidine-2'-phenylindole dihydrochloride (DAPI, 1:5,000 in 0.1 M PBS) for 30 s at room temperature, coverslipped using ProLong Gold Antifade Mountant (ThermoScientific) and dried for 48 h. Slides were stored at 4 °C in the dark.

**Immunohistochemistry**

To also examine c-Fos expression at the protein level, some mice were anesthetized with an overdose of sodium pentobarbital 90 min after the memory inference test, followed by transcardial perfusion with 20 mL of ice-cold 0.1 M PBS and 20 mL of ice-cold 4% PFA (pH 7.1 – 7.2). Brains were extracted, and post-fixated in 4% PFA (pH 7.1 – 7.2) at 4 °C overnight. Then brains were cryoprotected in 30% sucrose in 0.1 M PBS for 2-3 days at 4 °C. Forty micrometer-thick coronal sections were cut on a cryostat (Leica) and collected in 0.1 M PBS with 0.01% sodium azide. Sections were stored at 4 °C until later use.

Every fourth section of the dorsal hippocampus (AP: -1.46 mm to -2.30 mm, relative to Bregma) was selected for staining (four sections per animal). Sections were washed in 0.1 M PBS for 10 min and then permeabilized using 0.3% Triton-X in 0.1 M PBS for 5 min. Next, sections were rinsed in 2% Normal Donkey Serum (NDS, Jackson Immuno Research) and 0.3% Triton-X in 0.1 M PBS for 50 min to block non-specific antibody binding. Then, sections were washed in 0.1 M PBS 3 times for 5 min each. Afterwards, sections were incubated with primary antibodies against c-Fos (guinea pig anti-c-Fos, 1:1,000, Synaptic Systems #226-308) and glutamate decarboxylase 67 (GAD67, marker for GABAergic neurons; mouse anti-GAD67, 1:500, Merck #MAB5406), diluted in 2% NDS and 0.3% Triton-X in 0.1 M PBS overnight at room temperature. The next day, sections were washed 3 times in 0.1 M PBS for 5 min each, and incubated with fluorophore-conjugated secondary antibodies: donkey anti-guinea pig Alexa Fluor 647 (1:750; Invitrogen) and donkey anti-mouse Alexa Fluor 488 (1:800; Invitrogen), diluted in 2% NDS and 0.3% Triton-X in 0.1 M PBS for 2 h at room temperature. All procedures starting from the secondary antibody incubation onwards were performed in the dark. Subsequently, sections were incubated with DAPI (1:5,000) in 0.1 M PBS for 1 min, then mounted on glass slides, left to dry and coverslipped using Fluorsave mounting medium (Sigma-Aldrich). Slides were stored at 4 °C in the dark.

To examine $GABA_A$ receptor activity, 20-µm thick coronal sections of the dorsal hippocampus (AP: -1.46 mm to -2.30 mm, relative to Bregma) were made, and mounted directly on SuperFrost Plus slides. Sections were left to dry at -20 °C for 2 h, and post-fixed in 4% PFA (pH 7.1 – 7.2) for 75 min, followed by washing in 0.1 M Tris-HCl-Buffered Saline (TBS, pH 8) twice for 2 min each. Sections were stored at -70 °C for later use. Sections were stained with primary antibodies against c-Fos (guinea pig anti-c-Fos, 1:1,000, Synaptic Systems #226-308) and phospho-$GABA_A$ receptor (rabbit anti-phospho-GABA-RB, 1:100, Merck #SAB4503762). Secondary antibodies were donkey anti-guinea pig Alexa Fluor 647 (1:750) and donkey anti-rabbit Alexa Fluor 488 (1:800). TBS was used instead of PBS. All other procedures were similar as described before.

To examine c-Fos expression within noradrenergic neurons of the locus coeruleus (LC; AP: -5.34 mm to -5.68 mm, relative to Bregma), coronal sections of 40 µm were made and post-fixed in 4% PFA (pH 7.1 – 7.2) for 75 min, followed by washing twice in 0.1 M PBS for 2 min each.

Sections were stored in 0.1 M PBS with 0.01% sodium azide at 4°C for later use. Every second section of the LC was selected (four sections per animal), and incubated with primary antibodies against c-Fos (guinea pig anti-c-Fos, 1:1,000, Synaptic Systems #226-308) and tyrosine hydroxylase (mouse anti-TH, marker for noradrenergic and dopaminergic neurons, 1:300, Sigma-Aldrich #MAB318), diluted in 2% NDS and 0.3% Triton-X in 0.1 M PBS at room temperature overnight. Secondary antibodies were donkey anti-guinea pig Alexa Fluor 647 (1:750) and donkey anti-mouse Alexa Fluor 488 (1:800).

**Imaging and quantification**

Sections for both FISH and immunohistochemistry were imaged using the Axioobserver with Sample Finder AI from Zeiss at 20X magnification and a numerical aperture of 0.8. The hippocampus was subdivided into granule cell layer of the dentate gyrus (DG), pyramidal cell layer of CA3 (CA3py), pyramidal cell layer of CA1 (CA1py), striatum radiatum of CA3 (CA3sr) and striatum radiatum of CA1 (CA1sr). c-Fos activity and glutamatergic relative activity was examined in the pyramidal layers only, while GABAergic relative activity was determined in the striatum radiatum layers. These regions of interest were drawn manually according to the stereotaxic mouse brain atlas (Paxinos & Franklin, 2001).

For FISH, the number of fluorescent cells were counted using an automated segmentation pipeline (https://github.com/AngelosDid/Region-CellSegmentation). Briefly, QuPath (Bankhead et al., 2017) was trained by manually selecting fluorescent cells (approximately 8 cells per image, in each channel). The trained QuPath network was used by StarDist (Haghofer et al., 2023) to automatically identify neurons in the sections, separately for each region of interest. Detected cells were filtered out when the area was <0.0035 $mm^2$ and if their mean fluorescence intensity fell below a set threshold relative to the background. Finally, detection was manually corrected, if necessary, in ImageJ software. The number of *c-Fos*-positive cells (>3 *c-Fos* puncta) per $mm^2$ averaged across the four sections per animal was used as a measure of neuronal activity in the region of interest. Relative GABAergic activity was calculated by dividing the number of neurons showing *c-Fos-Slc32a1* overlap by the total number of *Slc32a1*-positive (>3 *Slc32a1* puncta) neurons, and subsequently averaged across the four sections per animal. Similarly, relative glutamatergic activity was calculated by dividing the number of neurons showing *c-Fos-Slc17a7* overlap by the total number of *Slc17a7*-positive (>3 *Slc17a7* puncta) neurons, and averaged across the four sections per animal. Neurons were said to overlap when >80% of their areas overlapped.

For immunohistochemistry, immunopositive cells were manually counted by a researcher blind to treatment, separately for each region of interest, normalized to the area (cells/$mm^2$), and averaged across four sections per animal. Further, the number of cells showing overlap between c-Fos and GAD67, c-Fos and GABAR-P or c-Fos and TH were counted manually (when >80% of the cells' area overlapped), and normalized to the total number of GAD67- cells, GABAR-P- or TH-positive cells. Relative GABAR-P activity was examined in the pyramidal layer of area CA3 only, and relative noradrenergic activity (i.e. c-Fos-TH overlap) was examined in the LC.

**Cannula placement verification**

Cannula placement was verified histologically using cresyl violet staining by an observer blind to the treatment. Mice with needle tips located outside the dorsal hippocampus or with extensive tissue damage were excluded from final analyses.

**Statistical analyses**

All statistical analyses were performed using JASP 0.16.3.0 (Jasp Team 2024).

The reward-seeking bias on the memory inference test was analyzed with one- or two-way ANOVA with systemic or intrahippocampal drug treatment as between-subjects parameter(s). One-sample *t*-tests were used to determine whether the reward-seeking bias was different from zero (i.e. chance level). Reward-seeking behavior for the two auditory cues separately on the memory inference test was analyzed with a repeated-measures ANOVA with drug treatment as between-subject parameter and cue (rewarded vs. non-rewarded or rewarded vs. novel) as within-subject parameter. When appropriate, post-hoc tests were used to determine the origin of the significance, and *P*-values were corrected for multiple testing using a Bonferroni correction.

For FISH and immunohistochemistry, the number of fluorescent neurons was analyzed with independent samples *t*-tests with drug treatment (i.e. CORT 10 mg/kg vs. vehicle or yohimbine 3 mg/kg vs. saline) as between-subject parameter for each hippocampal subregion separately. Two-way ANOVA was used to analyze the number of fluorescent neurons in the CORT-propranolol experiment with CORT and propranolol treatment as between-subject parameters. When appropriate, post-hoc tests were used to determine the origin of the significance. *P*-values were corrected for multiple testing using a Bonferroni correction.

Plasma CORT levels were log-transformed and one-way ANOVA was used to analyze differences between drug treatment groups (CORT 3 or 10 mg/kg vs. vehicle or yohimbine 1 or 3 mg/kg vs. saline). When appropriate, post-hoc tests were used to determine the origin of the significance, and *P*-values were corrected for multiple testing using a Bonferroni correction.

All statistical test were two-tailed, and $P < 0.05$ was accepted as statistical significance.
