## Supplemental Figure 1 for "Glucocorticoids impair memory inference through noradrenergic disruption of GABAergic regulation in hippocampal CA3"

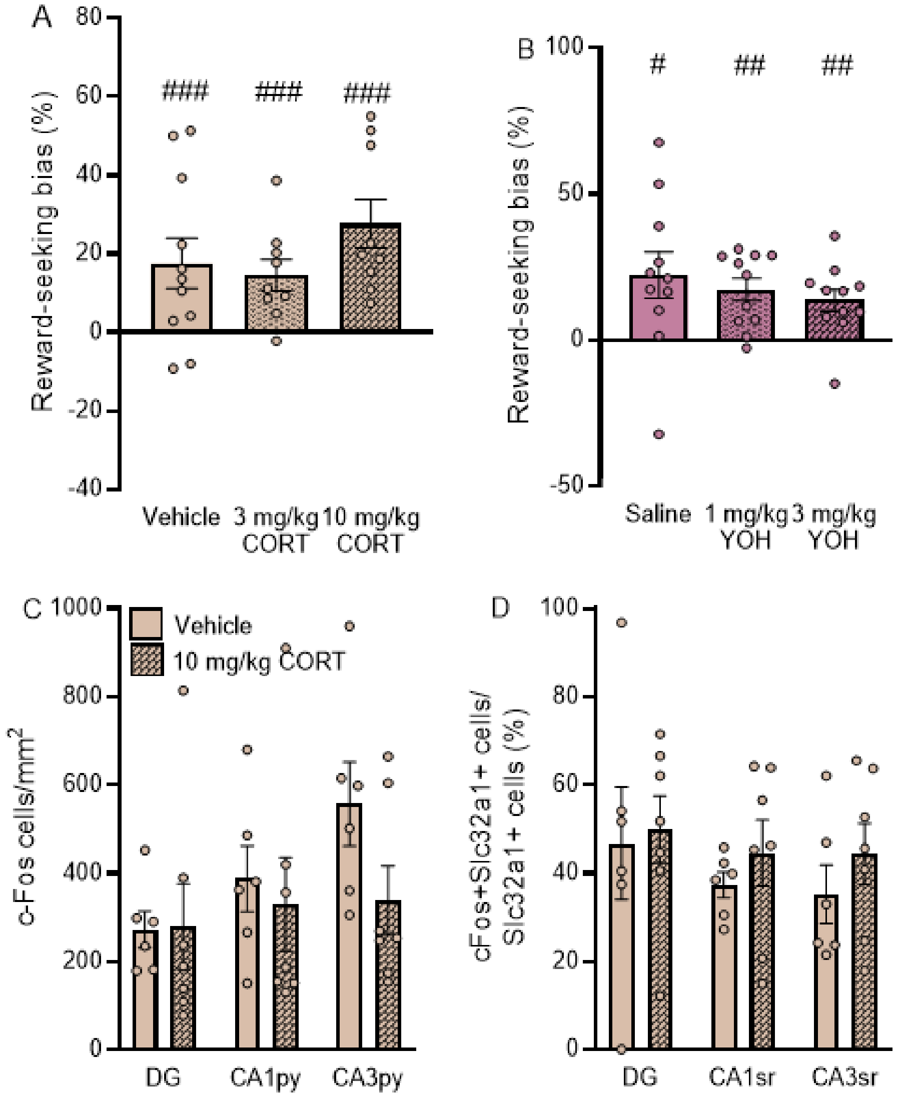


**Figure S1: Corticosterone and yohimbine did not affect the reward-seeking bias or hippocampal activity in animals re-exposed to the visual cues. A.** CORT did not affect the reward-seeking bias ($F_{2, 26}$ = 1.33, $P$ = 0.282), and all reward-seeking biases were greater than 0 (vehicle: $t_{10}$ = 6.27, $P$ < 0.001; 3 mg/kg CORT: $t_8$ = 6.40, $P$ < 0.001; 10 mg/kg CORT: $t_8$ = 11.17, $P$ < 0.001), in animals re-exposed to the visual cues. Vehicle: $n$ = 11, 3 mg/kg CORT: $n$ = 9, 10 mg/kg CORT: $n$ = 9. **B.** Yohimbine also did not affect the reward-seeking bias ($F_{2, 30}$ = 0.61, $P$ = 0.553), and all reward-seeking biases were greater than 0 (saline: $t_{10}$ = 2.79, $P$ = 0.019; 1 mg/kg YOH: $t_{10}$ = 4.51, $P$ = 0.001; 3 mg/kg YOH: $t_{10}$ = 3.56, $P$ = 0.005). Saline: $n$ = 11, 1 mg/kg YOH: $n$ = 11, 3 mg/kg YOH: $n$ = 11. **C.** CORT did not affect the number of neurons showing *c-Fos*-expression in the hippocampus (area CA1: $t_{11}$ = 0.43, $P$ = 0.673; CA3: $t_{11}$ = 1.79, $P$ = 0.101; DG: $t_{11}$ = 0.06, $P$ = 0.953) in animals exposed to the visual cues. **D.** CORT also did not affect *c-Fos*-expression in GABAergic cells in the hippocampus (CA1: $t_{11}$ = 0.84, $P$ = 0.419; CA3: $t_{11}$ = 0.95, $P$ = 0.362; DG: $t_{11}$ = 0.22, $P$ = 0.830) in animals exposed to the visual cues. Vehicle: $n$ = 8, 10 mg/kg CORT: $n$ = 7. Data represent mean ± SEM, circles represent individual data points. $^{\#}P$ < 0.05 vs. chance level, $^{\#\#}P$ < 0.01 vs. chance level, $^{\#\#\#}P$ < 0.001 vs. chance level.
