## Supplemental Figure 2 for "Glucocorticoids impair memory inference through noradrenergic disruption of GABAergic regulation in hippocampal CA3"

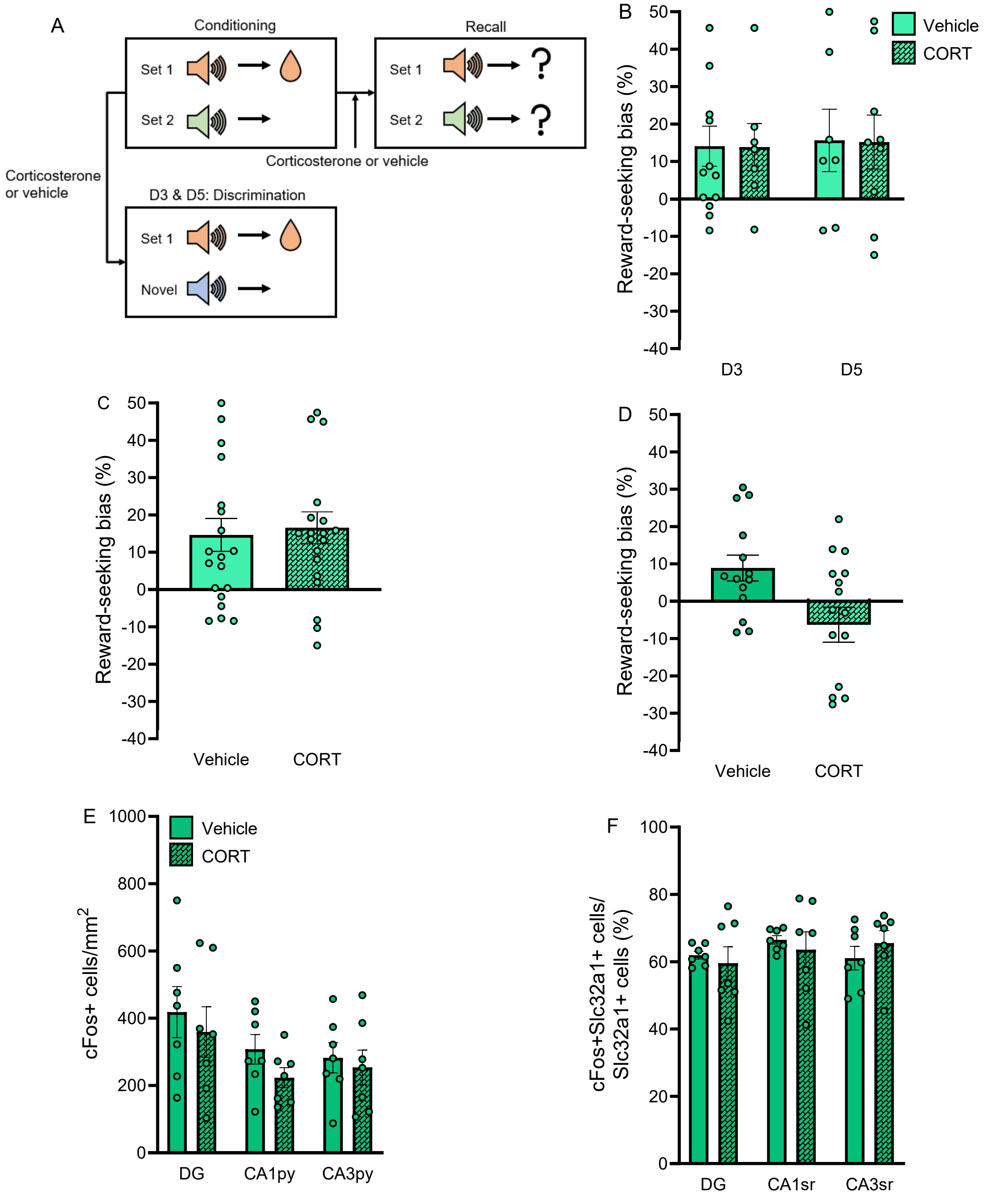


**Figure S2: Corticosterone did not affect animals' ability to discriminate between auditory cues, and CA3 activity was not associated with CORT effects on recall. A.** Experimental design of the discrimination and recall test. Animals were trained to associate one auditory cue (e.g. set 1) with a 15% sucrose reward. Additionally, on day 3 and 5 vehicle-control or 10 mg/kg CORT was administered (counterbalanced within the animal), and 45 min later animals were exposed to the rewarded auditory cue and a novel auditory cue. Animals were exposed to the rewarded and the novel cue 5 times each (discrimination test), and were subsequently trained in the conditioning stage as described before. Forty-five minutes prior to the recall test (24 hours after the last conditioning training), CORT (10 mg/kg) or vehicle-control was administered, and animals were re-exposed to the auditory cues. **B.** The reward-seeking bias did not depend on drug treatment (i.e. CORT or vehicle; $F_{2, 32} = 0.002$, $P = 0.961$), the day of treatment (i.e. day 3 or 5; $F_{2, 32} = 0.05$, $P = 0.833$), or their interaction ($F_{2, 32} < 0.001$, $P = 0.989$). **C.** The reward-seeking biases pooled between the 2 groups (independent of the day of treatment). **D.** CORT did not impair the retrieval of the auditory cue-reward association ($t_{27} = 1.91$, $P = 0.067$). Vehicle: $n = 14$, 10 mg/kg CORT: $n = 16$. **E.** CORT treatment did not affect *c-Fos*-expression in the recall task ($t_{27} = 0.42$, $P = 0.686$). **F.** CORT treatment did not affect *c-Fos*-expression in GABAergic cells in the recall task ($t_{27} = 0.88$, $P = 0.398$). Vehicle: $n = 7$, 10 mg/kg CORT: $n = 7$. Data represent mean ± SEM, circles represent individual data points. *$P < 0.05$ vs. vehicle-control, $^{\#}P < 0.01$ vs. chance-level.
