## Supplemental Figure 3 for "Glucocorticoids impair memory inference through noradrenergic disruption of GABAergic regulation in hippocampal CA3"

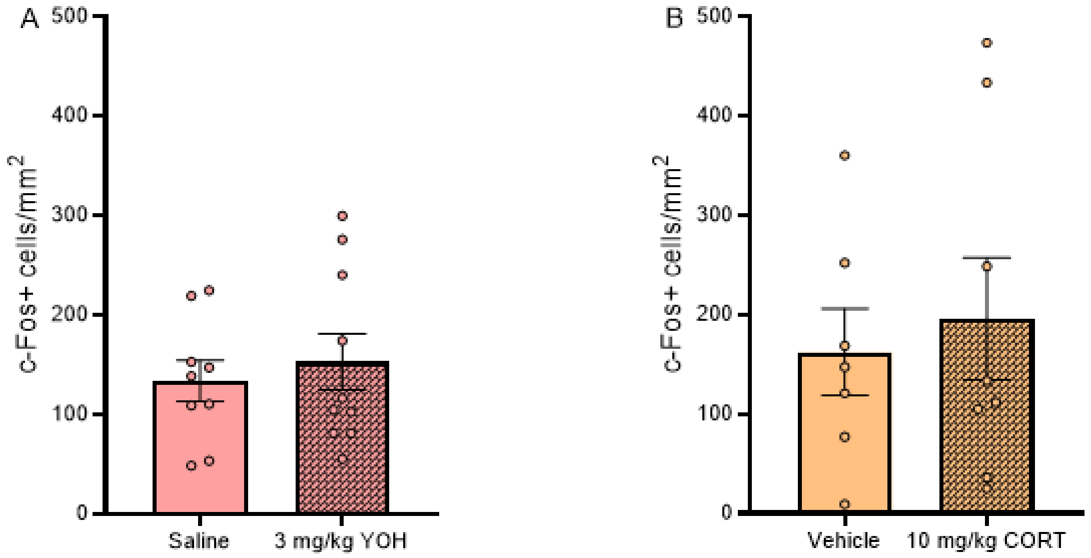


**Figure S3: Drug treatment did not alter c-Fos-expression in the LC. A**. Yohimbine treatment did not alter c-Fos-expression in the LC ($t_{17}$ = 0.17, *P* = 0.865). Saline: *n* = 9, 3 mg/kg YOH: *n* = 10. **B**. CORT treatment did not alter c-Fos-expression in the LC ($t_{22}$ = 0.04, *P* = 0.972). Vehicle: *n* = 10, 10 mg/kg CORT: *n* = 14. Data represent mean ± SEM, circles represent individual data points.
