## Supplemental Figure 4 for "Glucocorticoids impair memory inference through noradrenergic disruption of GABAergic regulation in hippocampal CA3"

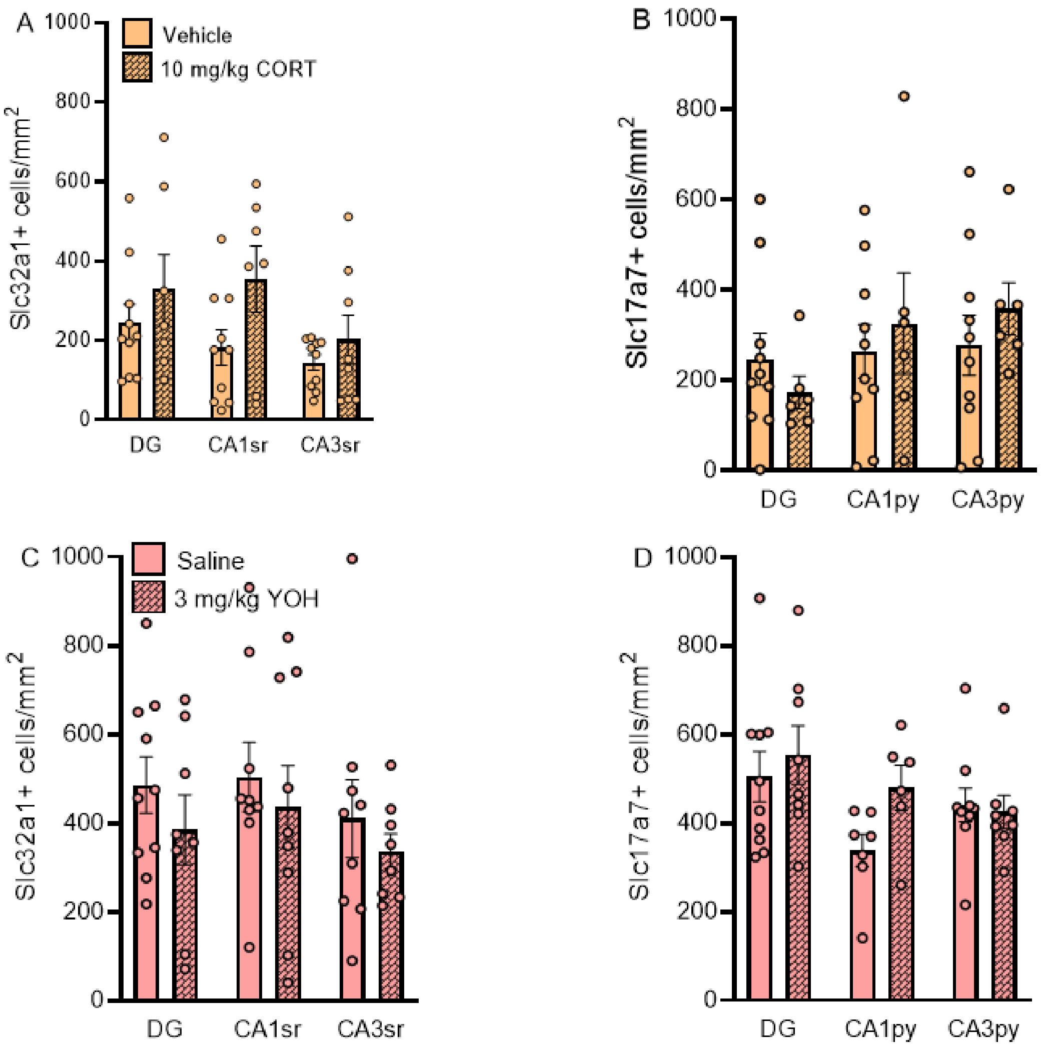


**Figure S4. Total *Slc32a1*- or *Scl17a7*-expression was not affected by drug treatment. A.** CORT treatment did not affect the total number of GABAergic cells in the hippocampus (CA1: $t_{16}$ = 1.97, $P$ = 0.201; CA3: $t_{16}$ = 0.98, $P$ = 0.343; DG: $t_{16}$ = 0.93, $P$ = 0.366). **B.** CORT treatment did not affect the total number of glutamatergic cells in the hippocampus (CA1: $t_{16}$ = 0.53, $P$ = 0.604; CA3: $t_{16}$ = 0.84, $P$ = 0.417; DG: $t_{16}$ = 0.92, $P$ = 0.373). Vehicle: $n$ = 9, 10 mg/kg CORT: $n$ = 9. **C.** Yohimbine treatment did not affect the total number of GABAergic cells in the hippocampus (CA1: $t_{17}$ = 0.56, $P$ = 0.585; CA3: $t_{17}$ = 0.74, $P$ = 0.473; DG: $t_{17}$ = 1.01, $P$ = 0.326). **D.** Yohimbine treatment did not affect the total number of glutamatergic cells in the hippocampus (CA1: $t_{17}$ = 2.29, $P$ = 0.129; CA3: $t_{17}$ = 0.30, $P$ = 0.767; DG: $t_{17}$ = 0.57, $P$ = 0.577). Saline: $n$ = 9, 3 mg/kg YOH: $n$ = 10. Data represent mean ± SEM, circles represent individual data points.
