## Supplemental Figure 5 for "Glucocorticoids impair memory inference through noradrenergic disruption of GABAergic regulation in hippocampal CA3"

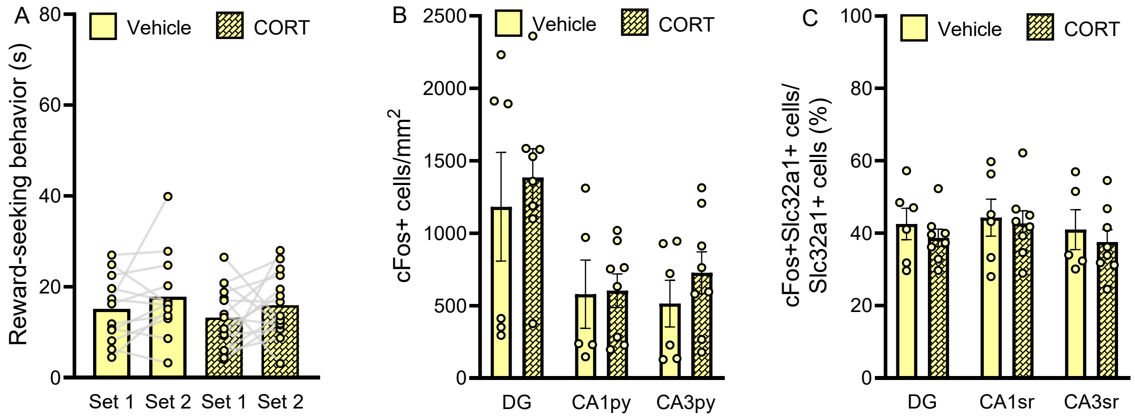


**Figure S5: When accuracy is no longer needed, CORT does not affect the CA3. A.** When both visual cues were rewarded during conditioning (both auditory cues may be inferentially linked to reward), CORT (10 mg/kg, systemically administered 45 min before the memory inference test) did not affect memory inference; animals showed similar reward-seeking behavior to the two auditory cues (main effect CORT: $F_{2,26}$ = 0.67, $P$ = 0.420; main effect cue: $F_{2,26}$ = 2.57, $P$ = 0.121; CORT × cue interaction: $F_{2,26}$ < 0.001, $P$ = 0.986,. Vehicle: $n$ = 13, 10 mg/kg CORT: $n$ = 15. **B.** No differences due to CORT-treatment in *c-Fos*-expression were found (CA1: $t_{12}$ = 0.10, $P$ = 0.921, CA3: $t_{12}$ = 0.98, $P$ = 0.346, DG: $t_{12}$ = 0.51, $P$ = 0.618). **C.** No differences due to CORT-treatment in *c-Fos*-expression in GABAergic cells were found (CA1: $t_{12}$ = 0.27, $P$ = 0.793, CA3: $t_{12}$ = 0.56, $P$ = 0.585, DG: $t_{12}$ = 0.83, $P$ = 0.423). Vehicle: $n$ = 6, 10 mg/kg CORT: $n$ = 8. Data represent mean ± SEM, circles represent individual data points.
