## Supplemental Figure 6 for "Glucocorticoids impair memory inference through noradrenergic disruption of GABAergic regulation in hippocampal CA3"

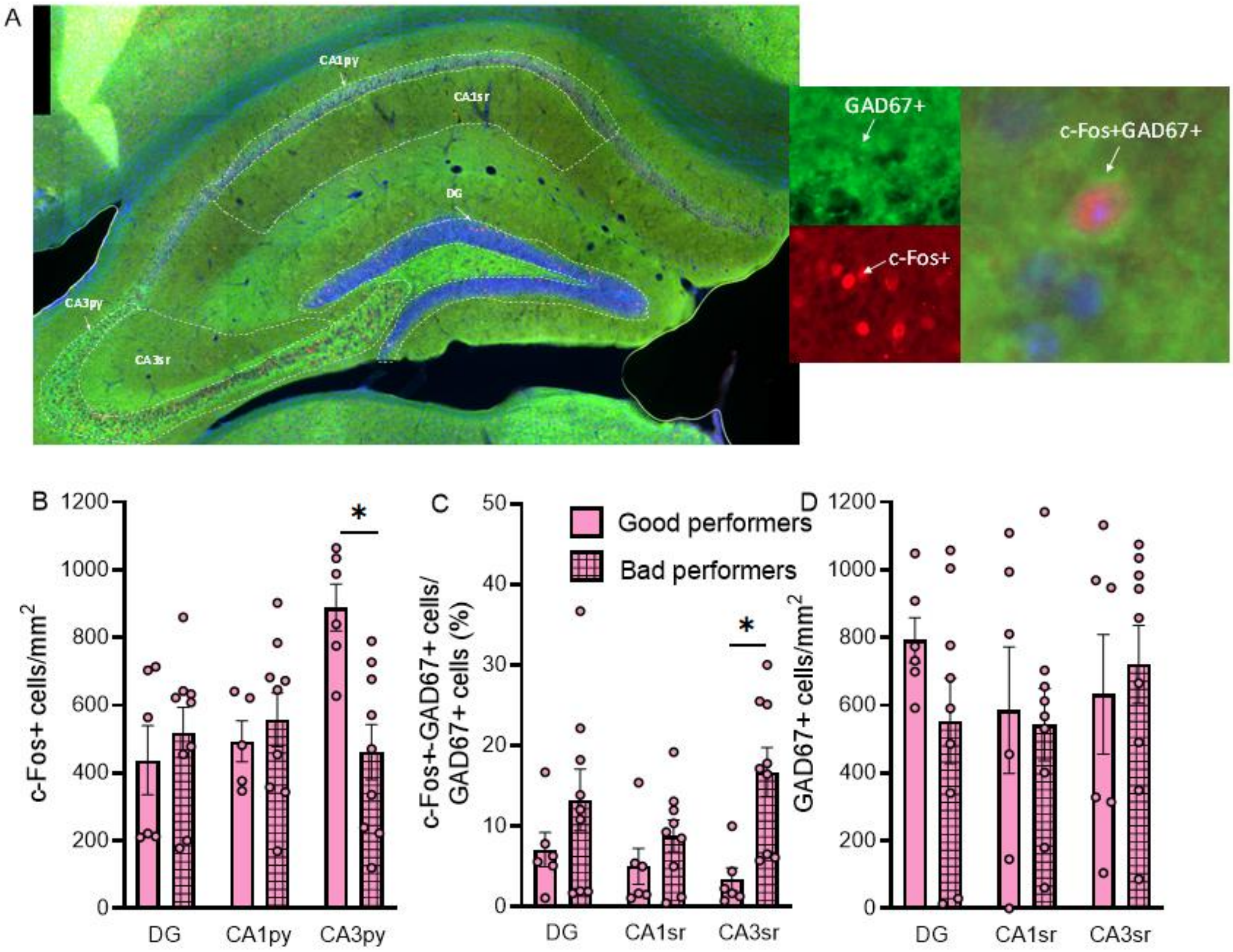


**Figure S6: A reduction in c-Fos-expression and increase in c-Fos-GAD67-co-expression in areas CA3 was associated with bad performance (reward-seeking bias <5%) on the memory inference test. A.** Images from an exemplary slice showing the hippocampal sub-regions and quality of immunohistochemistry staining (c-Fos+, GAD67+). **B.** Bad performance was associated with a reduction in the number of c-Fos-expressing cells in area CA3. CA3: $t_{13}$ = 3.71, $P$ = 0.009, CA1: $t_{13}$ = 0.54, $P$ = 0.599, DG: $t_{13}$ = 0.67, $P$ = 0.513. **C.** Bad performance was associated with an increased number of neurons showing c-Fos-GAD67-co-expression in area CA3. CA3: $t_{13}$ = 3.39, $P$ = 0.015, CA1: $t_{13}$ = 1.25, $P$ = 0.234, DG: $t_{13}$ = 1.23, $P$ = 0.242. **D.** Bad performance was not associated with a general increase in GAD67-expressing cells. CA3: $t_{13}$ = 0.47, $P$ = 0.647; CA1: $t_{13}$ = 0.46, $P$ = 0.650; DG: $t_{13}$ = 0.15, $P$ = 0.884. Good performers: $n$ = 6, bad performers: $n$ = 9. Data represent mean ± SEM, circles represent individual data points. *$P$ < 0.05 vs. good performance.
