## Supplemental Figure 7 for "Glucocorticoids impair memory inference through noradrenergic disruption of GABAergic regulation in hippocampal CA3"

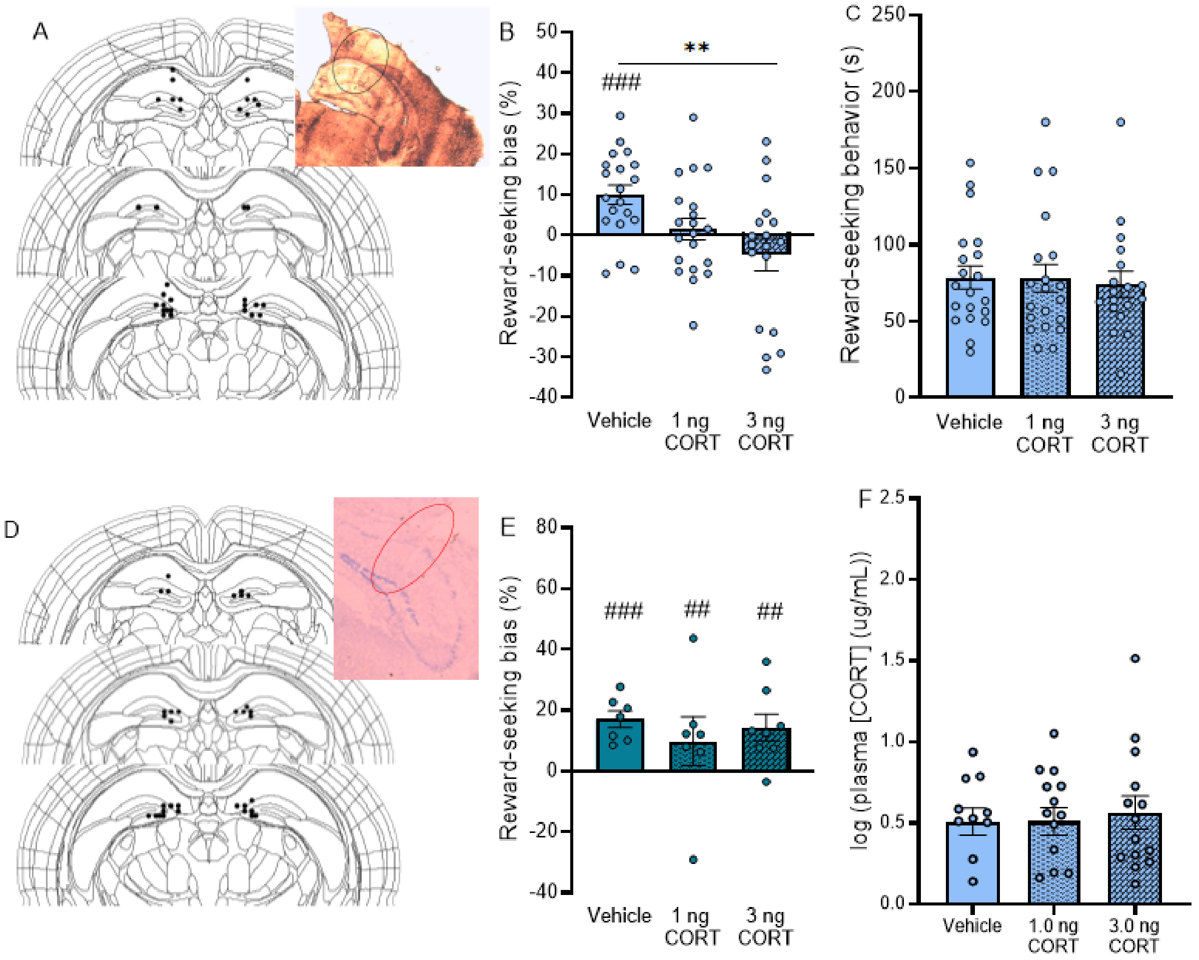


**Figure S7: Corticosterone administration into the hippocampus impaired the reward-seeking bias at 3 ng, and did not affect the reward-seeking bias when animals were re-exposed to the visual cues.** **A.** Histological image of an exemplary brain showing the position of the needles. The coordinates of the needle tips are also shown. **B.** 3.0 ng of CORT impaired the reward-seeking bias ($F_{2, 52} = 6.50$, $P = 0.003$; 1 ng: $t_{38} = 1.99$, $P = 0.157$; 3 ng: $t_{33} = 3.59$, $P = 0.002$, relative to vehicle). **C.** Total reward-seeking behavior was not affected by drug treatment ($F_{2, 52} = 0.49$, $P = 0.616$). Vehicle: $n = 21$, 1 ng CORT: $n = 21$, 3 ng CORT: $n = 17$. **D.** Histological image of an exemplary brain showing the position of the needles. The coordinates of the needle tips are also shown. **E.** CORT did not affect the reward-seeking bias when animals were re-exposed to the visual cues ($F_{2, 27} = 0.96$, $P = 0.650$), and all reward-seeking biases were greater than 0 (vehicle: $t_9 = 8.76$, $P < 0.001$; 1 ng CORT: $t_{10} = 4.43$, $P = 0.001$, 3 ng CORT: $t_8 = 3.55$, $P = 0.008$). Vehicle: $n = 10$, 1 ng CORT: $n = 11$, 3 ng CORT: $n = 9$. **F.** Treatment did not cause differences in CORT blood plasma levels. Vehicle: $n = 11$, 1 ng CORT: $n = 14$, 3 ng CORT: $n = 14$. Data represent mean ± SEM, circles represent individual data points. $^{**}P < 0.01$ vs. vehicle, $^{\#\#}P < 0.01$ vs. chance level, $^{\#\#\#}P < 0.001$ vs. chance level.
